# Antibiotic-free whole-cell biocatalytic fermentation: *Escherichia coli* with surface-displayed PETases for sustainable plastic degradation

**DOI:** 10.1101/2024.11.20.624590

**Authors:** Katherine Romero-Orejon, Hamid Reza Karbalaei-Heidari, Nediljko Budisa, David Levin

## Abstract

Plastic pollution has increasingly burdened the environment, driving the need for natural degradation platforms that utilize microbial enzymes to break plastics down into monomers. In this study, we introduce a novel approach using *Escherichia coli* as a fermentative, antibiotic-free whole-cell biocatalyst with surface-displayed, genomically integrated PETases for efficient plastic degradation. PETases, a class of esterases, catalyze the hydrolysis of polyethylene terephthalate (PET) into mono-2-hydroxyethyl terephthalate (MHET). Surface display of these enzymes was achieved via gene fusions with an N-terminal cysteine (Cys) triacylated anchor, mediated by the Braun lipoprotein (Lpp) signal peptide. To circumvent issues associated with plasmids, - such as genetic instability and reliance on antibiotics - we used a Type I-F CRISPR-associated transposase to insert the genes directly into specific *E. coli* genome sites. Proper enzyme display and activity on the *E. coli* surface were confirmed through enzyme activity tests, Western blotting, and flow cytometry, with cells retaining PET degradation ability over multiple generations. High-performance liquid chromatography (HPLC) analysis assessed degradation efficiency, identifying byproducts such as bis (2-hydroxyethyl) terephthalate and terephthalic acid. This study establishes a proof-of-concept for efficient plastic degradation using engineered bacteria as robust, sustainable, and genomically stable whole-cell biocatalysts, providing a promising platform for addressing plastic waste management.

**Graphical Abstract:** 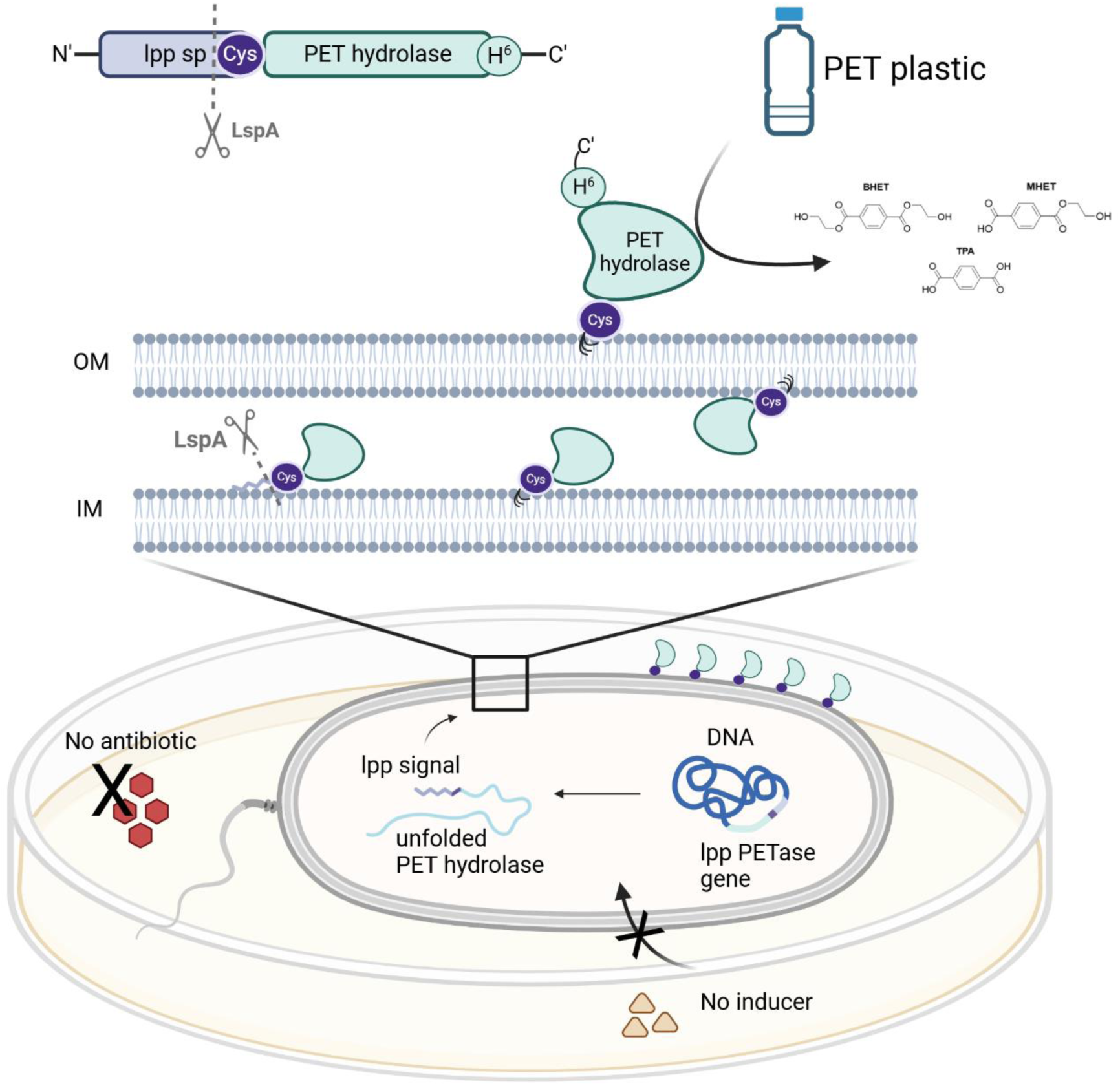

**One-sentence Abstract:** *Escherichia coli* was engineered as a fermentative, antibiotic-free whole-cell biocatalyst, featuring surface-displayed and genomically integrated PETases for efficient plastic degradation. This innovative approach has the potential to transform plastic recycling by enabling sustainable, large-scale degradation of plastic waste through environmentally friendly microbial systems.

## Introduction

Plastic pollution poses a serious threat to terrestrial and marine ecosystems, driving extensive biodiversity loss and ecosystem degradation, and making effective plastic waste management a critical global challenge. Each year around 400 million tonnes of plastic waste are generated worldwide, yet only about 9% is recycled^1,2^. Most plastic waste - roughly 80% - accumulates in landfills, while the rest is either incinerated or mismanaged, often leaking into the environment^2^.

Polyethylene terephthalate (PET), one of the “big five” plastics, along with polyethylene (PE), polypropylene (PP), polystyrene (PS), and polyvinyl chloride (PVC), is a widely used synthetic polymer made from ethylene glycol (EG) and terephthalic acid (TPA), commonly found in products like beverage containers and textile fibers, and accounts for approximately 18% of global polymer production^3^. Its extensive use has raised concerns about its long-term environmental and health impacts prompting increased research into effective removal or recycling strategies^4,5^. While physical and chemical recycling methods have been explored^6^, biological approaches are gaining traction as more sustainable and eco-friendly alternatives^7^.

In 2016, Yoshida and co-workers^8^ discovered *Ideonella sakaiensis* 201-F6, a bacterium from a PET bottle recycling facility in Japan, capable of efficiently degrading amorphous PET films. *I. sakaiensis* produces two key PET-degrading enzymes: PETase, a cutinase-like enzyme which breaks down PET into soluble intermediates: bis(2-hydroxyethyl) terephthalate (BHET), mono-(2-hydroxyethyl) terephthalate (MHET), and terephthalic acid (TPA); and MHETase, an esterase that further hydrolyses MHET into TPA and ethylene glycol (EG). These final products can either be recycled or used by the bacterium as a sole carbon source^8,9^.

Producing recombinant PET hydrolases using *Escherichia coli* presents a promising strategy for generating substantial quantities of enzymes to degrade PET plastic^10^. However, several challenges must be overcome to scale this method for industrial applications. In our view, despite advancements that enable isolated enzymes to degrade bulk plastic substrates within hours, they still face critical limitations: they are often unstable, especially over extended storage periods, and, most importantly, prohibitively expensive, making them unsuitable for large-scale industrial use^11,12^. Although enzyme immobilization is an appealing approach, it remains challenging due to difficulties in maintaining system stability and efficiency^13^.

Recent advancements in plastic biodegradation include innovative microbial co-display systems, exemplified by the work of Hu and Chen^10^. Their engineered *E. coli* strain, displaying FAST-PETase and the adhesive protein mfp-3 on its surface, degraded over 15% of amorphous PET within 24 hours at 30°C. Similarly, Gercke et al.^14^ embedded PETase in the outer membrane of *E. coli* using the inverse autotransporter YeeJ, achieving PET degradation five times more effectively at 25°C than free PETase at 30°C. The addition of rhamnolipids further enhanced this system, enabling 8% degradation of high-crystalline PET at an OD_600_ of 6 over three days at room temperature. Expanding on these efforts, Han et al.^15^ systematically studied the surface display of PET hydrolases, including FAST-PETase and MHETase, in *E. coli* to optimize PET hydrolysis. Their results showed that *E. coli* expressing pGSA-FAST-PETase at an OD_600_ of 1 achieved the highest hydrolytic activity, degrading 71.3% of PET powder in 24 hours at 50°C.

A high-throughput yeast surface display platform was developed to screen over 10⁷ enzyme mutants for PET film degradation^16^. In this system, enzymes were fused with the Aga2p protein, enabling multiple copies of each variant to be displayed on the *S. cerevisiae* cell membrane. The most effective variant identified, ICCG(H218Y), achieved 80% degradation of amorphous PET within 18 hours using 500 nM of the free enzyme. Additionally, a yeast-based system for displaying *Is*PETase enzymes was developed in *Pichia pastoris*^17^. In this approach, the GCW51 protein was used to anchor enzymes to the yeast cell wall, achieving 10% degradation of high-crystallinity PET within 18 hours at 30°C and pH 9.

*E. coli* BL21 (DE3)^18–21^, along with other prokaryotic and eukaryotic systems^22,23^, has proven effective for protein expression and whole-cell biocatalyst production in shaking-flask cultures. However, a significant limitation is their reliance on plasmid-based enzyme expression, which requires continuous supplementation of antibiotics and inducers and is prone to segregational instability, especially during fed-batch fermentation under the control of strong promoters. Over time, cells with reduced target gene expression can outcompete the engineered population, hindering large-scale production^24^. A more robust solution involves the chromosomal integration of PET-degrading enzymes under a constitutive promoter system, ensuring precise gene localization, controlled gene dosage, and improved genetic stability, all of which are critical for sustained industrial applications.

The Braun’s lipoprotein (Lpp) is a key outer membrane (OM) peptide protein that plays a crucial role in maintaining cell envelope integrity and supporting cell division. This lipoprotein is the most abundant protein in *E. coli*, having 58 residues in the mature form and directed to the outer membrane by a 20-amino acid signal peptide at the N-terminal^25,26^. Lpp exist in two forms: a “bound form” anchored to the OM of *E. coli* by three acyl group attached to an N-terminal cysteine residue and covalently linked after signal peptide cleavage while lysine is at the C-terminus to the peptidoglycan layer^26^. The “free-form” spans the OM and is exposed on the cell surface^27^. Lpp is initially synthesized in the cytoplasm as a pre-protein with a signal peptide, which directs its transport across the inner membrane (IM) through the SecYEG translocon^25^ (see Scheme 1). While tethered to the IM, two acyl chains are transferred to the sulfhydryl group of a conserved cysteine, catalyzed by diacylglyceryl transferase (Lgt). Signal peptidase II (LspA) then cleaves the signal peptide at the Gly20-Cys21 junction, leaving the acylated cysteine as the first residue. In γ-proteobacteria, like *E. coli*, this cysteine’s free amino group undergoes further acylation by apolipoprotein N-acyltransferase (Lnt), forming a mature lipoprotein with three acyl chains^28^. The Lpp signal sequence, containing the necessary motif, is recognized by the ABC transporter complex (LolCDE) which delivers it to the chaperone protein LolA, and subsequently to LolB at the OM. Naturally, Lpp’s lipid portion into the OM, while its C-terminal (Lys53) is covalently linked to the peptidoglycan layer via its ε-amino group^26^.

**Scheme 1.**
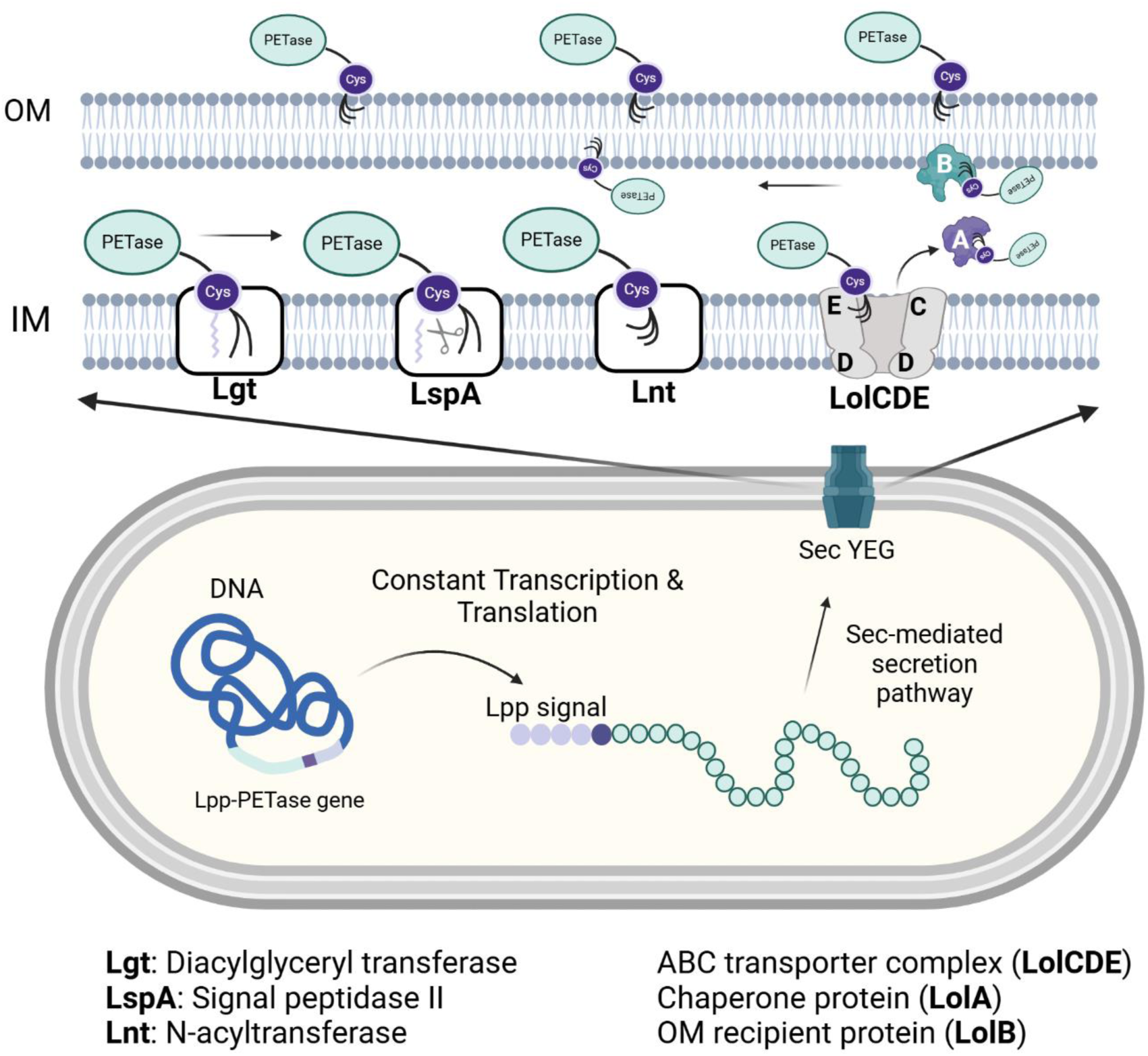
The proposed secretion pathway^25^ of Braun lipoproteins (Lpp) in *Escherichia coli*.

In our genetic construct, the C-terminal segment of the Lpp sequence was replaced with PET-hydrolyzing enzymes. Following the cleavage of the signal peptide in the Gly20-Cys21 junction, Cys21 undergoes triacylation, anchoring the PET hydrolases to the lipid bilayer at the N-terminus. By leveraging the Lpp promoter and the first 21 amino acids of the Lpp signal sequence, we drive lipidation and direct specific enzymes to the OM of *E. coli*. This strategy effectively targets PET-hydrolyzing enzymes to the *E. coli* OM while ensuring continuous expression of the downstream coding sequence, thereby enhancing enzyme expression, proper folding, localization, and activity.

However, maintaining stable enzyme expression over time, particularly in a system free from antibiotics and inducers, while achieving industrial scalability, presents a significant challenge. In this context, CRISPR (Clustered Regularly Interspaced Short Palindromic Repeats)-associated transposition technology is crucial for precise modification of bacterial genome, minimizing disruptions to existing coding sequences^29^. Specifically, Type I-F CRISPR-associated transposition (CAST) facilitates the efficient insertion of DNA payloads up to 30 kb with minimal off-target effects, making it ideal for developing stable, high-efficiency cell-based biocatalysts^30^. We employed a CRISPR-based transposon technology (VcTn6677), derived from *Vibrio cholerae* Tn6677 (*Vc*Tn6677)^31–33^. This approach enabled the integration of gene cassettes containing the constitutive Lpp promoter and Lpp signal peptide sequence upstream of PET-hydrolyzing genes into specific sites within the *E. coli* genome. This genome engineering strategy provides stable, continuous enzyme production and displays PET-hydrolyzing enzymes on the cell surface, eliminating the need for inducers or antibiotics.

In this study, our objective was to target PETase to the OM of *E. coli* by replacing the C-terminal part of the Lpp sequence with the PETase enzyme. This strategy aimed to anchor the enzyme on both the inner and outer surfaces of the OM, as well as in its free form, mimicking the natural behavior of Lpp^26^. We developed a plasmid-free *E. coli* chassis featuring a surface-displayed plastic degradation system. With chromosome-integrated enzymes, our platform provides a stable chassis that can be easily adapted to diverse plastic-degrading enzymatic systems. We also tackled critical challenges in selecting appropriate anchoring tags to maintain high protein expression while ensuring functional integrity. By eliminating genomic instability, the system operates without antibiotics, addressing sustainability, enzyme efficiency, and scalability - key challenges in designing a robust and fermentative platform for plastic degradation.

## Results

### Cys-PETase expression in *E. coli* BL21(DE3) and *E. coli* MS33 strains

To determine whether the native Lpp in *E. coli* interferes with Cys-PETase expression and proper localization, we generated an Lpp gene knockout strain of *E. coli* BL21(DE3) using CRISPR-Cas9-assisted genome editing method^30^. Amplification of the Lpp gene region with primers targeting the flanking regions in the *E. coli* MS33 genome confirmed the successful deletion of the gene fragment, resulting in the absence of mature Lpp protein expression (Figure S1). Next, we examined the overexpression of the pET29b-Lpp signal-PETase plasmid in both *E. coli* BL21 and *E. coli* MS33 (BL21-ΔLpp) at 16°C, 25°C, and 37°C. SDS-PAGE analysis revealed effective PETase expression in both strains at 16°C and 25°C (Figure S6). A comparison of expression at 4-hour and 18-hour intervals (Figure S3) showed that the intensity of the Cys-PETase band increased with longer expression times.

To confirm the successful display of Cys-PETase on the cell surface and to better understand its cellular localization, an immunoblotting assay was performed. The C-terminal His-tag of PETase was probed with an anti-His antibody across various fractions of IPTG-induced cells, revealing fluorescent signals in the 25 to 35 kDa range. These results confirmed that the Cys-PETase hybrid was successfully sorted to the OM in both strains (Figure 1a). However, some Cys-PETase was also detected in the IM and as a soluble fraction, highlighting the dynamic nature of protein translation and sorting.

**Figure 1.**
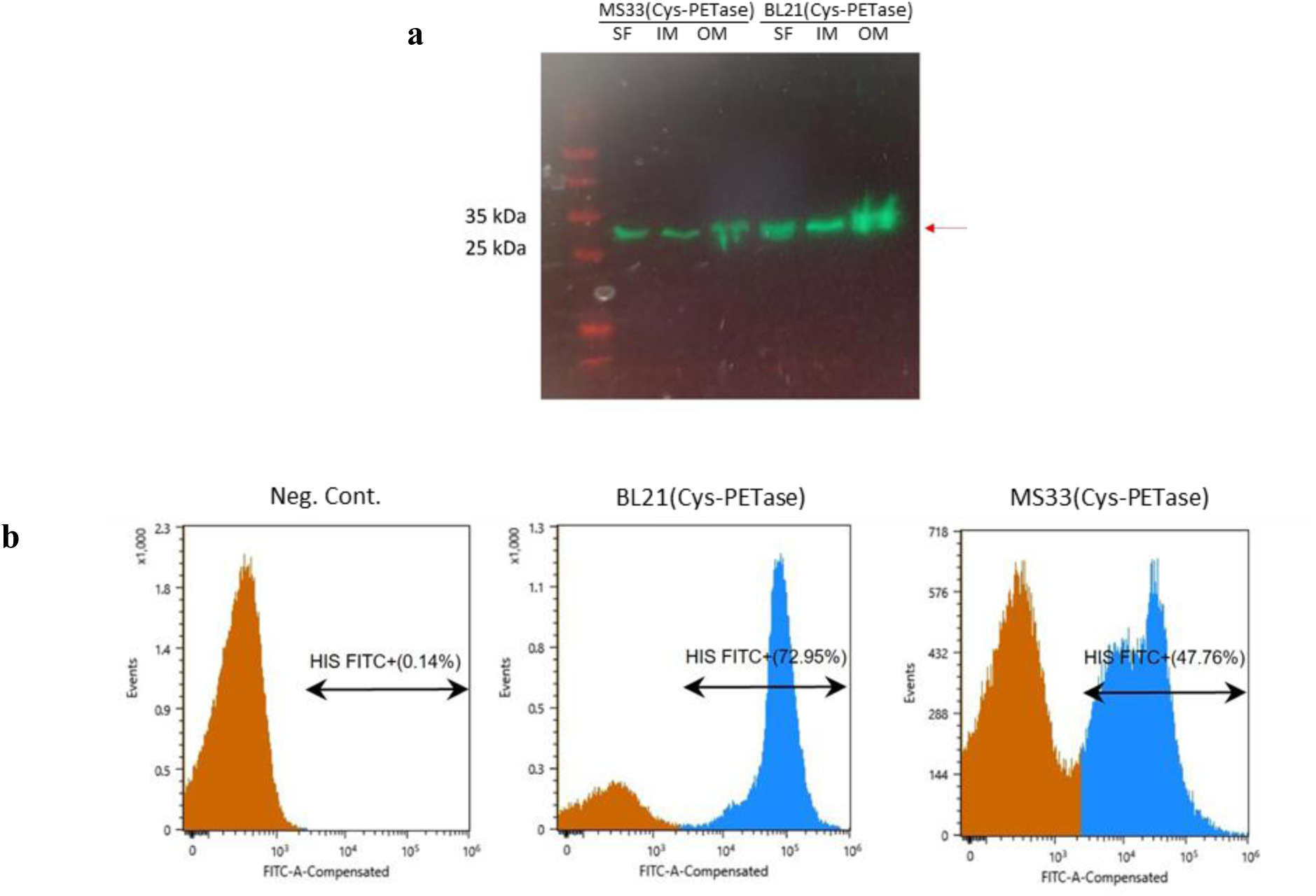
Surface display of Cys-PETase in *E. coli* BL21 and *E. coli* MS33. **a**) Western blotting of membrane fractions of *E. coli* BL21(Cys-PETase) and *E. coli* MS33(Cys-PETase). The outer membranes of both strains exhibited intense bands at ∼30 kDa, confirming the membrane-anchored enzyme production of the construct with N-terminal Lpp. In addition, the bands in the inner membrane and soluble fractions highlighted the dynamic nature of protein translation. L: ladder, SF: soluble fraction, IM: inner-membrane, OM: outer-membrane. **b**) The surface exposure of Cys-PETase was determined using flow cytometry analysis. *E. coli* BL21(Cys-PETase) exhibited a fluorescent signal of 72.95%, while *E. coli* MS33(Cys-PETase) showed a signal of 47.76%, confirming the surface exposure of the enzymes. *E. coli* BL21 without the plasmid served as a negative control, displaying a signal of only 0.14%. Samples were treated with an anti-His-Tag monoclonal antibody (6G2A9) and FITC-conjugated THE™.

Flow cytometry analysis (Figure 1b) revealed strong fluorescent signals (FITC emission) when the anti-His tag antibody interacted with Cys-PETase displayed on the surfaces of both the *E. coli* BL21 and *E. coli* MS33 strains, confirming the enzyme’s successful surface exposure. In the *E. coli* BL21 strain, 73% of the cells exhibited antibody binding upon Cys-PETase expression, while in the *E. coli* MS33 strain, this percentage was notably lower at 48%. As expected, the *E. coli* BL21 strain without the plasmid, which was used as a negative control, showed no fluorescent signal.

After confirming Cys-PETase expression and localization in both strains, esterase activity assays were performed to quantify the enzyme’s functional activity and evaluate the efficiency of its expression and surface display in each strain. Cys-PETase production was tested at three different temperatures (16°C, 25°C, and 37°C) by culturing both strains and assessing esterase activity using p-Nitrophenol Alkanoates (*p-*NPAlk) with varying alkyl chain lengths. One unit of *p*-NP esterase activity (U) is equivalent to 0.027 µM.min^-1^, being the specific activity 0.69 U/OD. As shown in Figure **2**a, the *E. coli* BL21(Cys-PETase) and *E. coli* MS33(Cys-PETase) strains exhibited significantly higher activity against *p*-NPAc (C2), particularly when cultured at 16°C and 25°C, with relative initial activities of 0.582 U/OD and 0.685 U/OD. Notably, the *E. coli* MS33(Cys-PETase) strain retained 85% of the activity of the *E. coli* BL21 strain under both temperature conditions (16°C and 25°C).

**Figure 2.**
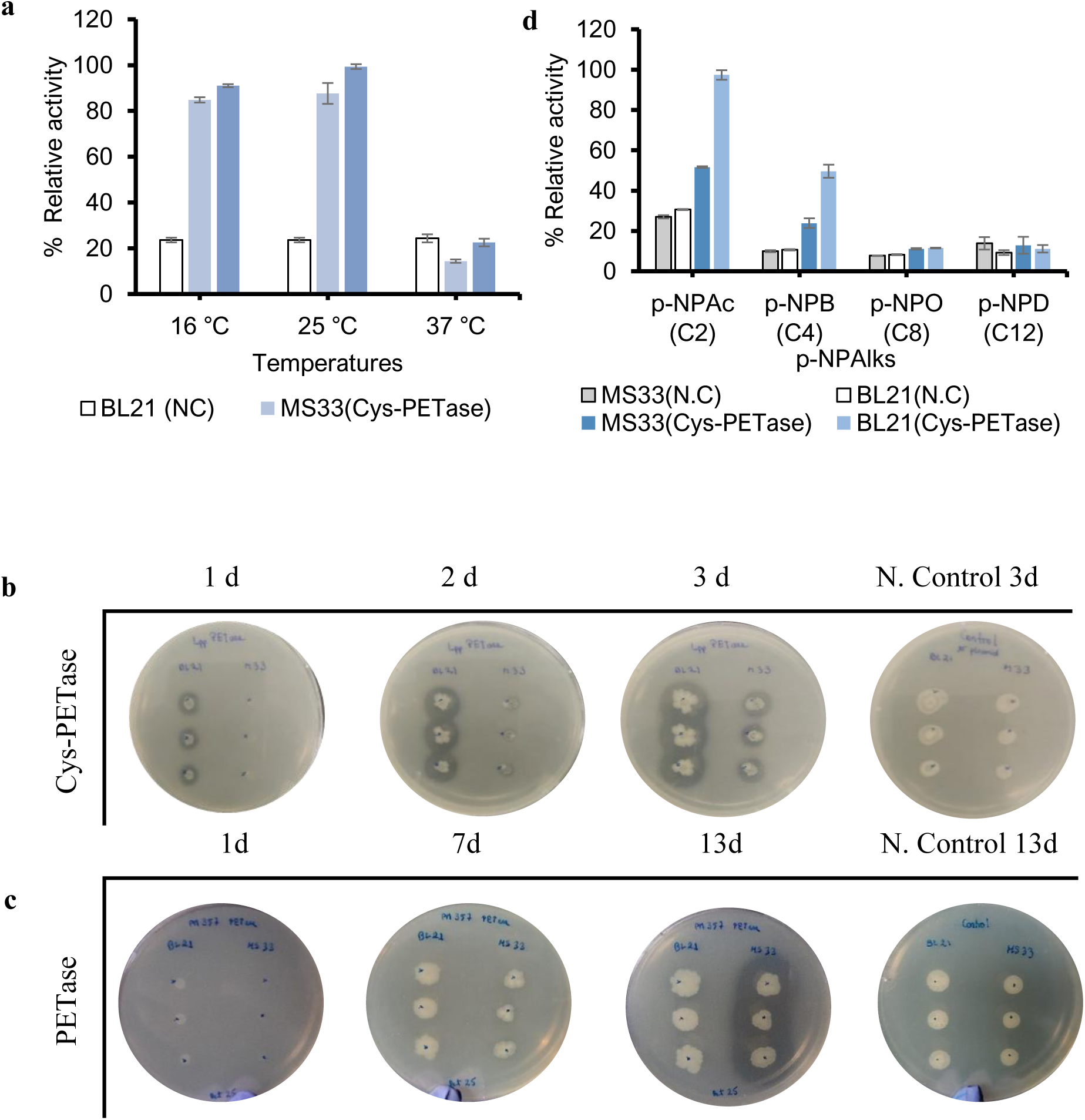
Optimization of the surface-displayed Cys-PETase on *E. coli* BL21 and *E. coli* MS33. **a**) The esterase activity assay against *p*-nitrophenyl acetate for surface-displayed enzymes, expressed at 16°C, 25°C, and 37°C, revealed that *E. coli* BL21(Cys-PETase) expressed at 25°C exhibited the highest relative activity, corresponding to 0.69 U/OD. **b**) The comparison of halo formation on BHET-LB agar plates between *E. coli* BL21 (Cys-PETase) and *E. coli* MS33 (Cys-PETase) demonstrated that *E. coli* BL21 (Cys-PETase) formed visible halos after one day, whereas *E. coli* MS33 (Cys-PETase) exhibited slower BHET hydrolysis. **c**) Cell membrane permeability was assessed through halo formation on BHET-LB agar plates. Strains expressing the PETase gene without the Lpp signal peptide were inoculated on the plates. Only *E. coli* MS33(PETase) hydrolyzed BHET over 13 days, suggesting enzyme leakage. All samples and controls were tested in triplicate. **d)** Esterase activity assay against p-Nitrophenyl Alkanoates (*p*-NPAlks) with different side chain lengths.: *p*-Nitrophenyl Acetate (C2), *p*-Nitrophenyl Butyrate (C4) *p*-Nitrophenyl Octanoate (C8) and *p*-Nitrophenyl Dodecanoate (C12).

The difference in Cys-PETase activity expressed in *E. coli* BL21 and *E. coli* MS33 was also visualized in BHET-LB agar plates. Consistent with the above data, the BHET-degrading activity of *E. coli* BL21(Cys-PETase) was superior to *E. coli* MS33(Cys-PETase) (Figure 2b), as indicated by the presence of halos around the colonies after one day of growth. In the case of *E. coli* MS33(Cys-PETase), its activity became noticeable only on the second day. The negative controls, *E. coli* BL21 and *E. coli* MS33 without plasmids, did not show any observable consumption of the BHET (Figure S4).

Overall, the results showed that knocking out the Lpp gene compromised cell envelope integrity, leading to reduced growth and decreased tolerance to low DMSO concentrations (used as a solvent for BHET in preparing BHET-LB agar plates). Additionally, cell growth at 37°C resulted in minimal esterase activity, likely due to inclusion body formation, highlighting the need for a slower translation rate to ensure proper enzyme targeting and folding in the periplasm, particularly with the presence of a disulfide bond. These findings identified 25°C as the optimal temperature for producing active, properly folded, and membrane-exposed Cys-PETase.

The PETase gene, lacking the Lpp signal peptide, was expressed in both strains and inoculated on BHET-LB agar plates to evaluate whether hydrolytic activity dependent on Lpp functionality or was related to membrane permeability and enzyme leakage in the *E. coli* BL21 or *E. coli* MS33 strains. As shown in Figure 2c, the *E. coli* BL21 strain expressing intracellular PETase, BL21(PETase), did not form halos around its colonies over 13 days of growth. In contrast, the *E. coli* MS33(PETase) strain produced a halo after 7 days, which intensified by day 13 (Figure S5). These results suggest that the Lpp knockout in *E. coli* MS33 compromised membrane stability, leading to BHET consumption due to the release of PETases a free cytoplasmic enzyme. PETase expression in *E. coli* MS33 was further confirmed by SDS-PAGE (Figure S6).

This findings align with earlier studies demonstrating that Lpp plays a mechanical role in maintaining cell envelope stability, with Lpp knockout strains showing softer membranes^34–36^. This confirms that PETase was successfully displayed on the surface of the *E. coli* BL21 membrane due to the Lpp signal peptide, making *E. coli* BL21 the preferred host to harboring the pET29b-Lpp-PETase-His plasmid. Additionally, the expressed Cys-PETase demonstrated stronger activity toward shorter alkyl chains, such as *p*-NPAc (C2), compared to longer chains like *p*-NPD (C12) (Figure **2**d). This observation is consistent with previous reports indicating PETase’s lower affinity for p-nitrophenol-linked longer aliphatic esters^8^, reinforcing its specificity compared to other cutinases or esterases.

### Activity measurement using PET particles as substrate

To evaluate the potential of PETase-displayed cells as efficient whole-cell biocatalysts, we quantified the products of PET hydrolysis - specifically, bis(2-Hydroxyethyl) terephthalate (BHET), mono-(2-hydroxyethyl) terephthalate (MHET), and terephthalate (TPA) - using high-performance liquid chromatography (HPLC). *Ideonella sakaiensis* PETase (*Is*PETase) catalyzes the degradation of PET, primarily yielding MHET, with BHET and TPA as secondary products^37^. BHET undergoes further hydrolysis to produce ethylene glycol (EG) and MHET, while MHET is subsequently hydrolyzed by *Is*MHETase, an essential enzyme, to produce TPA and EG^38^. These simpler organic molecules have potential as carbon sources for bacterial growth and may future applications in single-cell protein production and biotechnological applications.

As shown in Figure 3a, *E. coli* BL21(Cys-PETase) and *E. coli* MS33(Cys-PETase) demonstrated BHET hydrolyzing activity using a cell suspension of OD_600_ = 0.05 in the presence of 1 mM BHET, however, *E. coli* MS33(Cys-PETase) exhibited lower activity compared to *E. coli* BL21(Cys-PETase). Over 90 minutes, 15% of the BHET was consumed (blue), with MHET released as a product (gray column) (Figure 3b), confirming Cys-PETase functionality. In another experiment, BHET biodegradation was conducted at higher cell densities (OD_600_ = 3.0 and OD_600_ = 6.0), resulting in complete BHET consumption within 1 hour (Figure 3c). Additionally, a whole cell biocatalysis assay was performed on semi-crystalline PET powder applying different optical densities (OD_600_) ranging from 0.5 to 6.0. A release of 6 μM MHET was observed using an OD_600_ of 6.0 after 24 hours of incubation (Figure 3d – Figure S7) indicating enzyme activity inhibition when using PET particles with high crystallinity.

**Figure 3.**
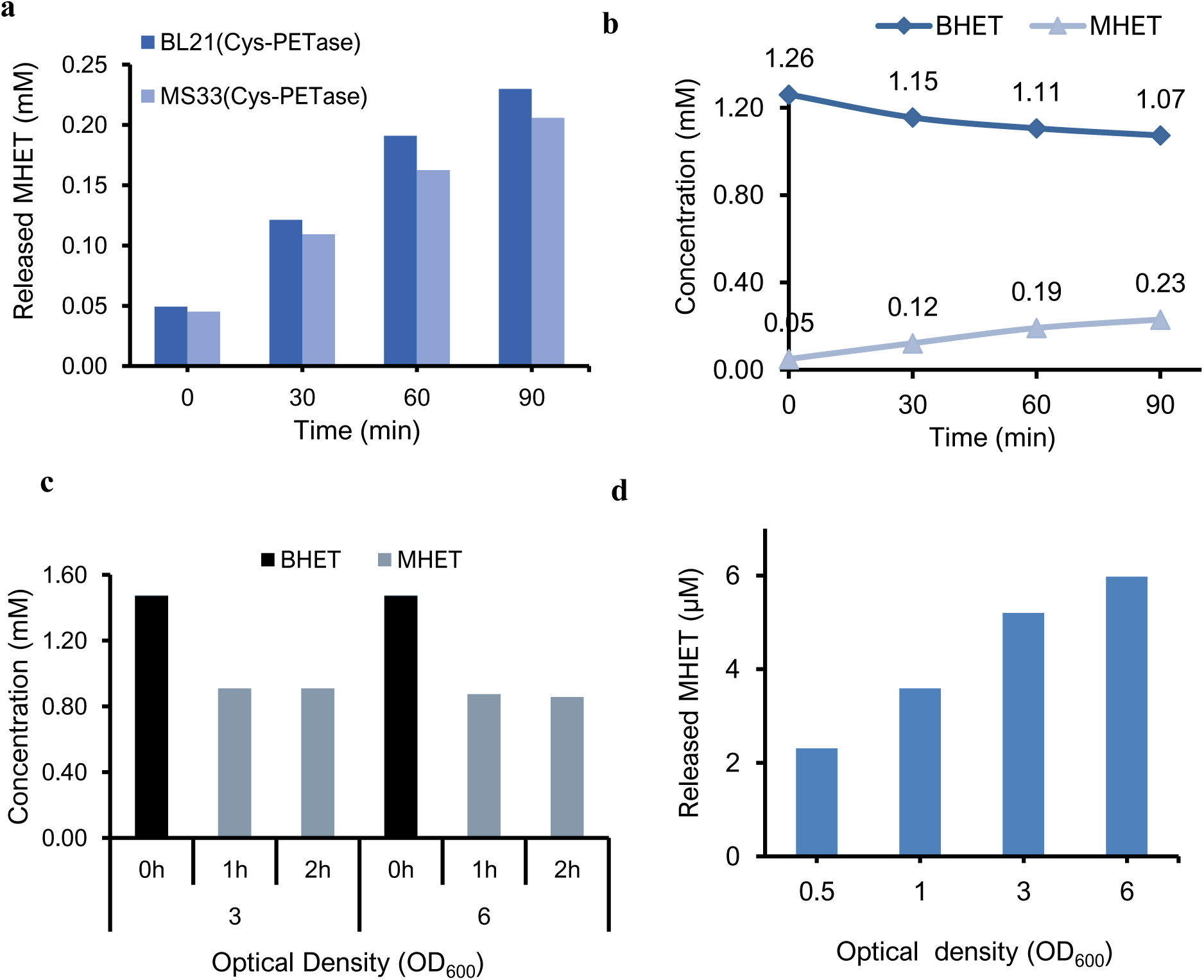
Biocatalysis of real substrates by surface-displayed Cys-PETase in *E. coli*. **a**) Comparison of BHET hydrolysis between *E. coli* BL21 (Cys-PETase) and *E. coli* MS33 (Cys-PETase) over time, using an initial OD_600_ of 0.05, **b**) BHET consumption (blue) and MHET released (gray) by *E. coli* BL21(Cys-PETase), the samples were analyzed every 30 minutes, **c**) BHET degradation at different optical densities (OD_600_) of *E. coli* BL21(Cys-PETase) revealed complete BHET hydrolysis within 1 hour at OD_600_ = 3. **d**) PET powder degradation by *E. coli* BL21(Cys-PETase) at different OD_600_. The graph illustrates the correlation between cell density of *E. coli* BL21(Cys-PETase) and MHET release after 24 hours of incubation at 37°C. The released product was quantified by High-performance liquid chromatography-photodiode array detector (HPLC-PDA).

### Genomically integrated Cys-PET hydrolases in *E. coli*

After confirming the effectiveness of the Cys-PETase strategy and demonstrating that the enzyme retained its functionality when anchored to the outer membrane (OM) of *E. coli* via the Lpp signal peptide, we expanded this approach to include additional PET hydrolases, such as MHETase^38^ and FAST-PETase^39^, and developed an antibiotic- and inducer-free system. We constructed donor plasmids with a constitutive Lpp promoter to eliminate the need for inducers: pDo-Lpp-Cys-PETase-His (PM530), pDo-Lpp-Cys-PETase-Cys-MHETase (PM532) as a polycistronic construct, and pDo-Lpp-Cys-FAST-PETase-His (PM570) (see Figure S8 for plasmid maps). Before applying these donor plasmids in mini-Tn transposition experiments (CAST) for site-specific DNA cassette genome integration, they were first transformed into *E. coli* BL21, and expression was confirmed in the plasmid-based system with appropriate antibiotic at 25°C. We then evaluated the whole-cell biocatalytic activity against BHET, confirming that the Cys-PET hydrolases were properly folded and actively degrading BHET.

As shown in Figure 4a, *E. coli* BL21(PM570) expressing Cys-FAST-PETase completely degraded BHET within 24 hours using a cell suspension at an OD_600_ of 0.1, demonstrating a faster reaction rate compared to the *E. coli* BL21(PM530) expressing Cys-PETase cells. As expected, FAST-PETase, which includes five mutations (S121E, D186H, R224Q, N233K, R280A)^39^, exhibited superior PET-hydrolytic activity compared to the wild-type PETase. Additionally, whole-cell biocatalysts *E. coli* BL21(PM530) and *E. coli* BL21(PM532), expressing Cys-PETase-Cys-MHETase, reduced BHET by approximately 50% over the same period, while the released MHET concentration was monitored (Figure 4b). Figure 4c shows that *E. coli* BL21(PM532) cells further converted released MHET into TPA, confirming the functionality of Cys-MHETase as a membrane-anchored enzyme. This was further supported by increased TPA production in *E. coli* BL21(PM532) cell’s reaction and by observing the degradation of MHET to TPA over time (Figure S9). Additionally, PET powder degradation was noted in Cys-FAST-PETase-expressing cells, which produced 28 µM of MHET and TPA, whereas Cys-PETase-Cys-MHETase-expressing cells (PM532) produced only TPA (Figure 4d).

**Figure 4.**
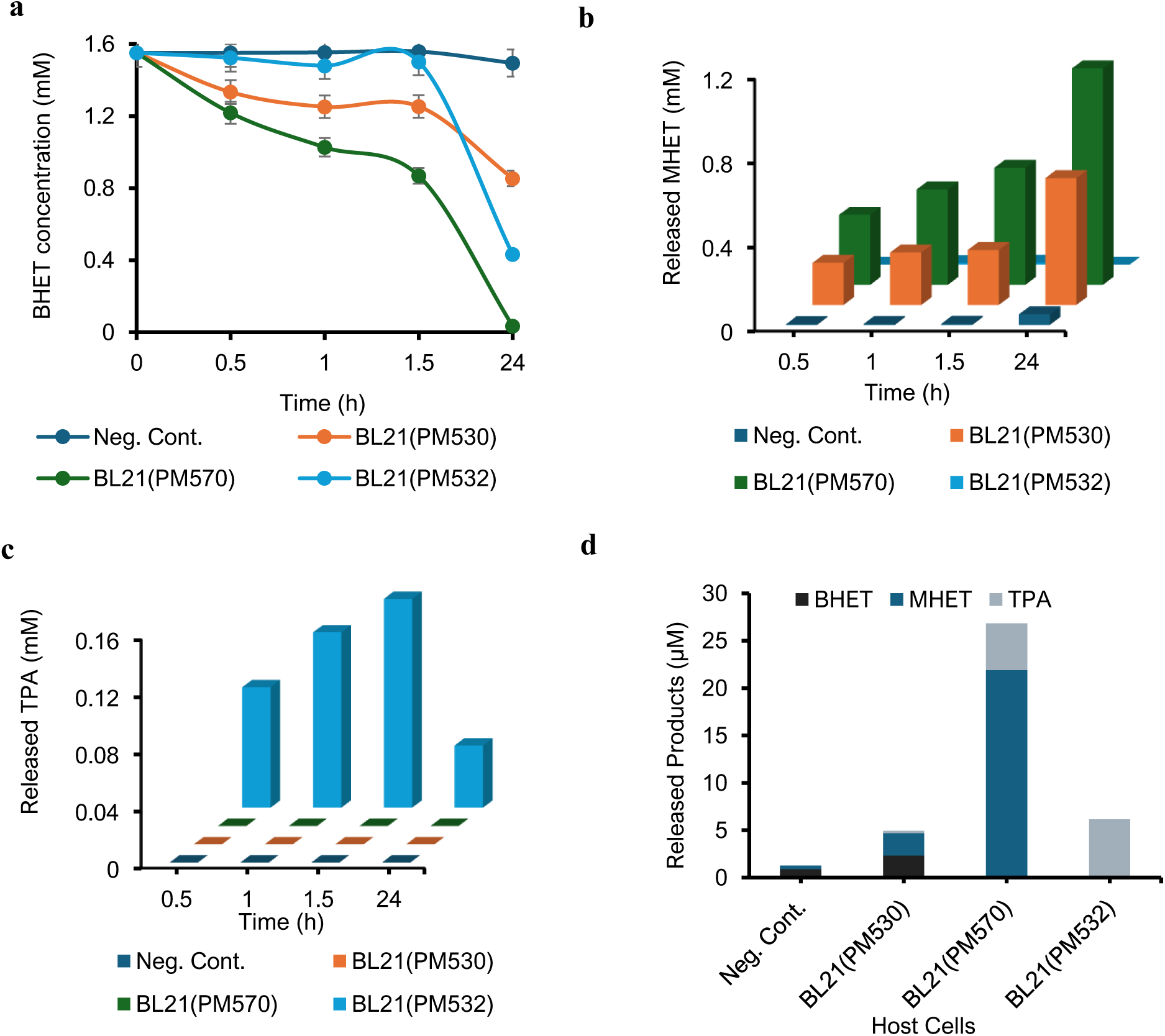
Comparison of product released during BHET and PET degradation using various Whole-Cell Biocatalysts. The biocatalysts include E. *coli* BL21(PM530) pDo-Lpp-PETase-His, *E. coli* BL21(PM570) pDo-Lpp-FAST-PETase-His and *E. coli* BL21(PM532) pDo-Lpp-PETase-Lpp-MHETase. **a)** BHET consumption, **b)** MHET released, **c)** TPA released over time, and **d)** Released products during PET powder hydrolysis.

Cells co-expressing two PET-hydrolyzing enzymes did not leave any MHET in the reaction mixture, unlike cells expressing Cys-PETase alone. However, they produced a lower TPA concentration compared to FAST-PETase expressing cells. This suggests that some of the products may have been metabolized by the cells or lost as an insoluble fraction during product extraction for HPLC analysis.

The next step involved integrating the gene cassettes into the *E. coli* genome using CRISPR-associated transposases (CAST). *E. coli* BL21(DE3) expression cells were co-transformed with the pEffector-array (PM452 or PM511) and one of the plasmids -PM530, PM532, or PM570. Mini-Tn transposition was then initiated using 5 µg/mL of anhydrotetracycline to induce the expression of the multi-complex protein transposition machinery, facilitating precise site-specific DNA integration without introducing double-strand breaks. Colony PCR analysis of the induced and isolated transformants confirmed the successful integration of the Cys-PETase and Cys-FAST-PETase gene cassette under the constitutive Lpp promoter. The cassettes were inserted 49 bp downstream of the *kdgA* gene sequence in the *E. coli* MS75 strain and downstream of the *marR* gene in the *E. coli* MS92 strain, respectively (Figure S10).

The genomically integrated strains, each harboring a single copy of the Lpp-PET hydrolase gene cassette, were cultured in LB medium without inducers or antibiotics. The *E. coli* MS75 strain was incubated at 25°C for various time intervals, and Western blot analysis was performed to evaluate the expression and surface localization of Cys-PET hydrolases. As shown in Figure 5a, a strong green signal from the anti-His tag antibody, at approximately 30 kDa was detected in the soluble fraction, inner membrane (IM), and outer membrane (OM) lanes of the *E. coli* MS75 cell pellet. In contrast, *E. coli* BL21 cells carrying the pET29-PETase plasmid without the Lpp signal, used as a negative control, exhibited a signal only in the soluble fraction, with no detection in the IM or OM fractions. These results confirm the successful expression of Cys-PETase and its localization in both the IM and OM of the *E. coli* MS75 strain.

**Figure 5.**
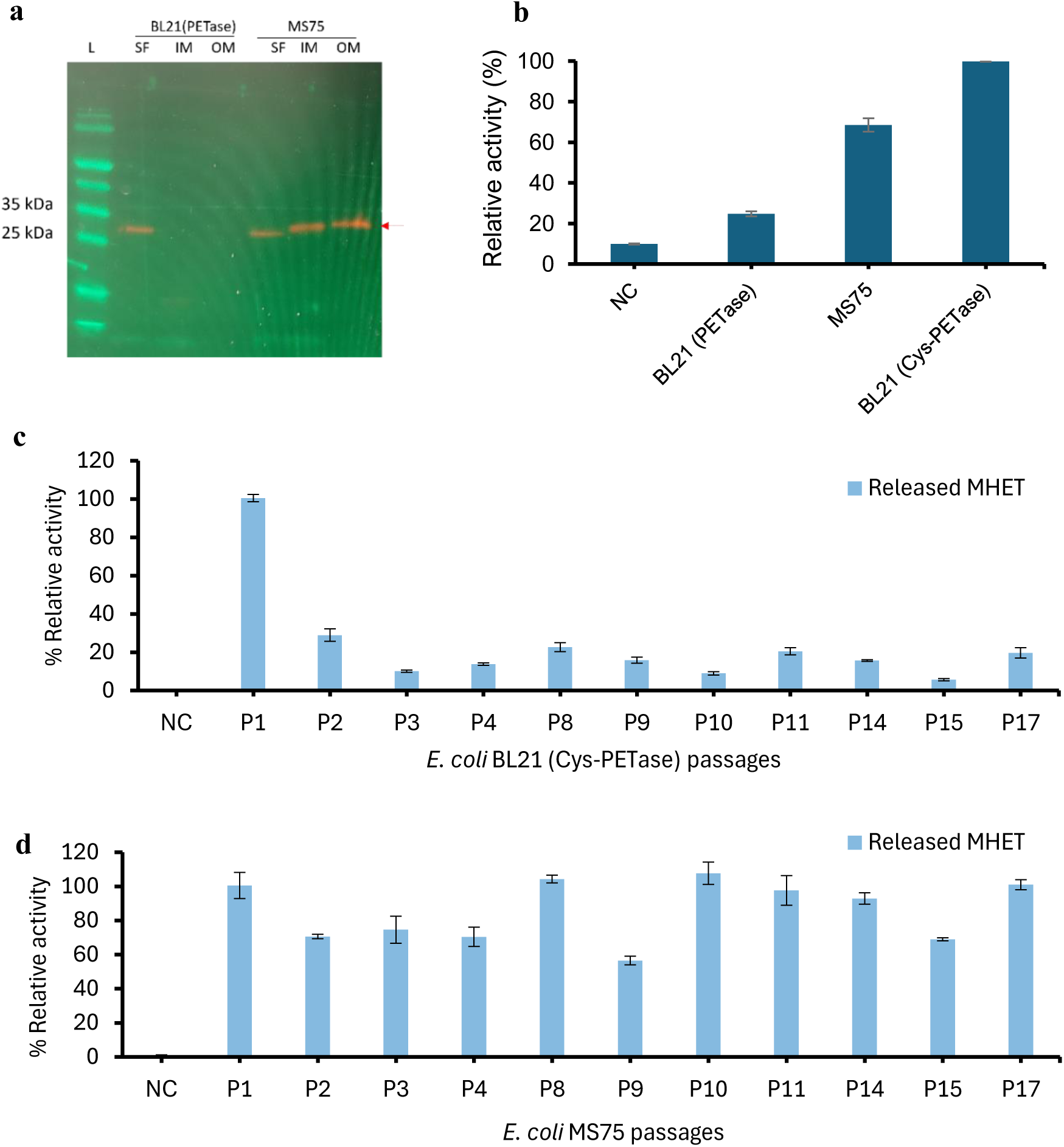
Surface-displayed Cys-PETase in *E. coli* MS75. **a**) Western blotting of *E. coli* BL21(PETase) and *E. coli* MS75 cells. Bands at ∼30 kDa were observed in both the IM and OM fractions of the *E. coli* MS75, while *E. coli* BL21(PETase) without Lpp signal showed a band exclusively in the soluble fraction, confirming the membrane-anchored enzyme production in *E. coli* MS75. L: ladder, SF: soluble fraction, IM: inner membrane, OM: outer membrane. **b)** Comparison of the esterase activity between *E. coli* BL21(PETase), *E. coli* BL21(Cys-PETase) and *E. coli* MS75 against *p*-NPAc (C2). Negative control (NC): no cells with *p*-NPAc. **c**) Comparison of the whole-cell biocatalytic activity against BHET by *E. coli* BL21(Cys-PETase) and **d)** *E. coli* MS75 over 17 passages. NC: BHET incubated without cells.

Esterase activity in the *E. coli* MS75 strain was further validated using a chromogenic substrate (Figure 5b), with the strain exhibiting 70% of the activity observed in plasmid-based Cys-PETase cultures. This reduced activity is likely due to the lower gene dosage, or fewer gene copies, in the *E. coli* MS75 strain compared to the plasmid-bearing cells. As the surface exposure of PETase via Lpp anchoring involves a gradual maturation process, sufficient incubation time is required to achieve optimal enzyme localization. To evaluate the impact of incubation time on maximum enzyme display, incubation periods from 1 to 3 days were tested. The results revealed that the highest level of enzyme exposure was reached after 2 days of incubation at 25°C, with activity remaining stable over this period (Figure S11). Furthermore, halos observed around *E. coli* MS75 colonies on BHET-LB agar plates (Figure S12) confirmed that the genomically integrated PETase is effectively surface-localized on the OM and functionally active.

### Stable expression of Cys-PETases in genomically integrated strains in an antibiotic- and inducer-free system

To assess the long-term stability of Cys-PETase expression, the engineered *E. coli* MS75 strain was compared to *E. coli* BL21 (PM447: T7 p-Cys-PETase) through serial passages in culture (Figure 5c &d). While the plasmid-based Cys-PETase exhibited higher initial activity in the first passage (Figure 5c), its relative activity declined rapidly, dropping to approximately 40% by the second passage (P2) and below 20% after 17 passages. In contrast, the *E. coli* MS75 strain retained over 90% of its initial activity by the seventeenth passage (P17), with only minor fluctuations observed, reaching a minimum of 60% of initial activity at P9 Figure 5d). These results indicate that plasmid-based systems are prone to genetic instability or attenuation of expression during extended culturing, likely due to the metabolic burden on the cells. By comparison, the *E. coli* MS75 strain demonstrated superior stability, maintaining consistent expression of the integrated PETase gene over time in an antibiotic- and inducer-free environment, ensuring sustained enzyme production.

The *E. coli* MS92 strain (Cys-FAST-PETase integrated strain) demonstrated a significantly higher rate of BHET degradation over a 24-hour period (Figure 6a) and produced HPLC-detectable amounts of MHET and TPA as final products when incubated *E. coli* MS92 OD_600_=1 with PET particles, despite their >50% of crystallinity (Figure 6b). Halos formed around *E. coli* MS92 colonies on BHET-LB agar plates confirmed that the genomically integrated Cys-FAST-PETase is not only functional but also more active than that of the *E. coli* MS75 strain (Figure 6c). Remarkably, *E. coli* MS92 maintained its activity against BHET across 10 passages (Figure 6d). To address high crystallinity PET film was pretreated with 10 M NaOH for 17 hours to reduce crystallinity^40^. After one week pf incubation with the *E. coli* MS92 strain, SEM images showed significant degradation (Figure 6e). The treated PET film exhibited a scratched surface, whereas the control PET film without cell incubation remained intact, serving as a negative control.

**Figure 6.**
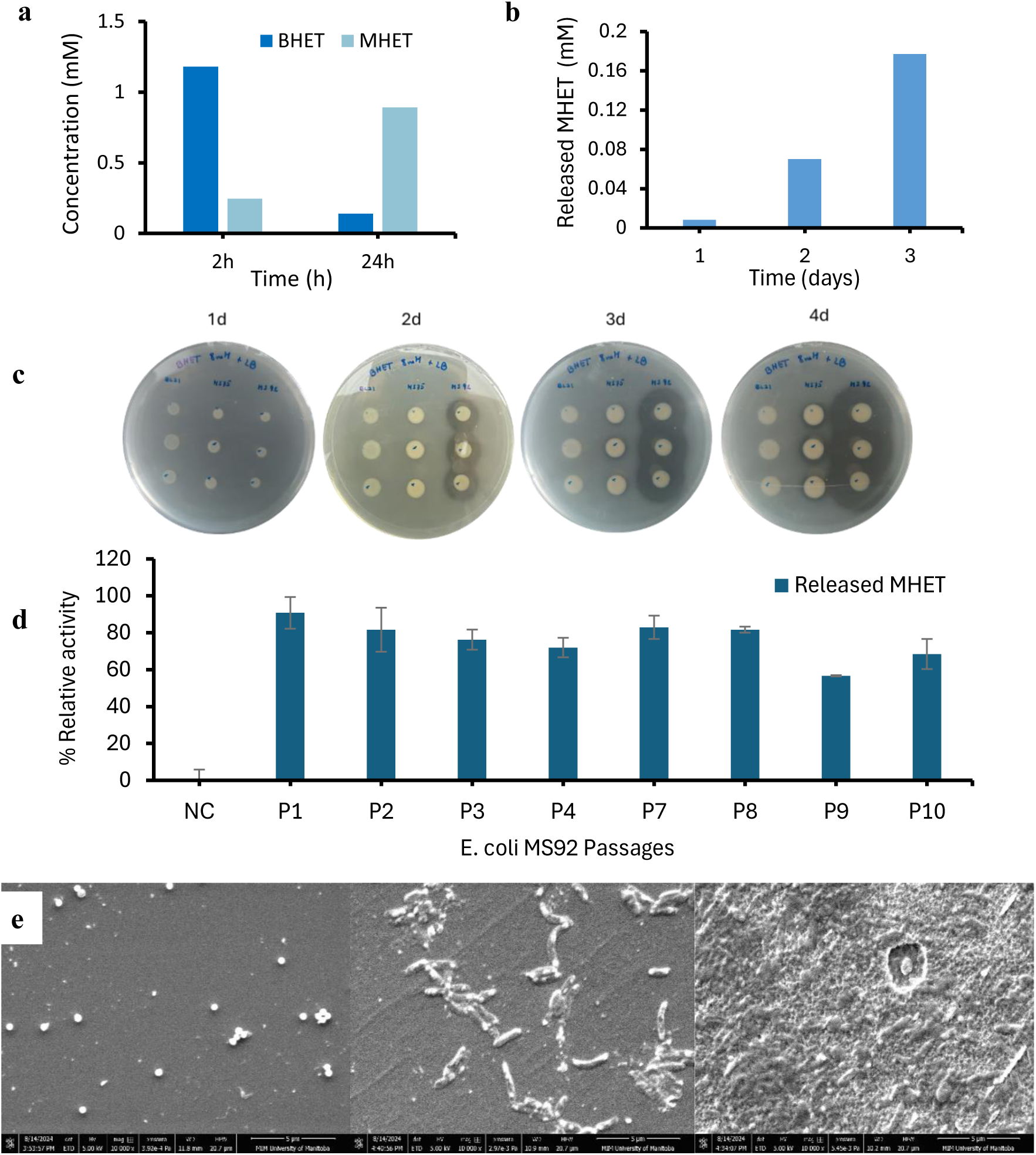
Assessment of the activity of genomically integrated *E. coli* MS75 (Cys-PETase) and *E. coli* MS92 (Cys-FAST-PETase). **a)** BHET hydrolysis by *E. coli* MS92 after 24 hours at OD_600_ = 1.0, **b**) PET powder biodegradation by *E. coli* MS92 (OD_600_ = 1), showing MHET released over 3 days, **c)** Comparison of halo formation on BHET-LB agar plates between *E. coli* MS75 and *E. coli* MS92, **d)** Comparison of whole-cell biocatalytic activity against BHET by *E. coli* MS92 over 10 passages. Cys-FAST-PETase expression in *E. coli* MS92 remained stable after 10 passages. NC: BHET incubated without cells. **e**) SEM images of commercial PET film before and after incubation with *E. coli* MS92 cells (scale bar: 5µm). Left: PET film after incubation with buffer, middle: *E. coli* MS72 attached to PET film, right: PET film after incubation with *E. coli* MS72.

## Discussion

We have employed Synthetic Biology to develop a whole-cell biocatalyst that immobilizes and simultaneously exposes the enzymes PETase, FAST-PETase, and MHETase to the bulk solvent using an N-terminal cysteine (Cys) triacylated anchor. This strategy eliminates the need for enzyme purification and prevents aggregation, which often reduces catalytic activity. Furthermore, by utilizing advanced genetic editing tools like CRISPR-associated transposase, we enhanced these biocatalysts by removing the dependency on inducers and antibiotics, significantly lowering production costs.

The Lpp signal peptide was selected to anchor PETases to the outer membrane (OM) of *E. coli* due to its abundance and its ability to target proteins to the cell surface^26^. This signal peptide directs the enzyme to the OM and provides a cysteine anchor for immobilization. Our results demonstrated that *E. coli* BL21 cells expressing Cys-PETase, Cys-MHETase, and Cys-FAST-PETase exhibited activity against both chromogenic compounds and real substrates such as BHET and PET powder. Notably, *E. coli* BL21(Cys-FAST-PETase) produced up to 25 µM of released product, compared to only 5 µM with *E. coli* BL21(Cys-PETase). This aligns with previous findings, as FAST-PETase is a mutated version of PETase with enhanced activity^39^.

A similar anchoring strategy for displaying PETase was reported by Heyde et al.^19^ and Zhang et al.^21^, both utilizing the LppOmpA fusion protein. Heyde et al. developed a screening platform to detect PET-degrading enzymes, while Zhang et al. enhanced adhesive interactions with material surfaces by genetically incorporating 3,4-Dihydroxy-L-phenylalanine (DOPA) at specific, solvent-exposed residues within the linker region of *E. coli*’s outer membrane protein A (OmpA). In both studies, protein expression was controlled by the rhamnose-inducible rhaPBAD promoter, and semi-crystalline PET powder was used to assess proper enzyme folding. Product formation was monitored by measuring absorbance at 240 nm. However, direct comparisons between the studies were difficult due to variations in optical density (OD_600_) values and incubation times. Future research could explore the relative effectiveness of different anchoring strategies under standardized conditions using the same PETases.

We used an N-terminal fusion construct to anchor PETases in the outer membrane (OM), with the PETase positioned at the C-terminus yielding the most optimal configuration. Proper folding of PETase requires stable disulfide bonds^11,41^, making the redox environment critical for its functionality. This approach proved particularly effective for signal peptide-free expression of PETase, as the reducing conditions of the cytosol often destabilize disulfide bonds, causing them to revert to free thiol groups. A similar outcome was observed using the LppOmpA anchor, which involved fusing the C-terminus of LppOmpA to a single-domain camelid-derived antibody fragment (nanobody; NB) with high affinity for GFP, subsequently linked to the N-terminus of the target protein^19^.

Expression temperature played a critical role in achieving proper enzyme folding and signal peptide maturation, likely due to improved coordination between translation and secretion rates, which directly impacts enzymatic activity. Testing three temperatures revealed that 25°C was optimal for esterase activity in both *E. coli* BL21(Cys-PETase) and the genomically integrated *E. coli* MS75 and *E. coli* MS92 strains. Similar expression temperatures (25–30°C) have also been employed in other studies using AIDA^10^, INPNC^10^, LppOmpA^19^ and pGSA^15^ as anchors. The choice of anchoring protein is crucial for ensuring a homogeneous population of bacterial cells displaying the target enzyme on the surface^42^. Furthermore, operating at room temperature offers a potential advantage in designing a cost-effective process that integrates PET degradation with product conversion.

*E. coli* BL21(Cys-PETase) demonstrated the ability to degrade semi-crystalline PET powder at an OD_600_ of 6, with degradation showing a strong linear correlation to enzyme concentration, enabling the use of high cell densities. Similarly, YeeJ-PETase exhibited a linear degradation rate up to an OD_600_ of 6; however, activity stagnated between OD_600_ values of 6 and 8, and activity declined at OD_600_ values above 10^14^. In contrast, PETase displayed on the surface of yeast (*Pichia pastoris*)^17^, was evaluated for PET degradation, but was effective only at very low concentrations. This inhibition at high enzyme concentrations is likely due to a molecular crowding effect, as recently documented in enzymatic PET depolymerization studies^43^.

Recent advancements highlight the use of microbial cell chassis, such as *E. coli* and yeast, to display plastic-degrading enzymes like PETases on their surfaces. Various anchors, including Lpp-OmpA, AIDA, and the inverse autotransporter YeeJ, have emerged as promising strategies for PET degradation in large-scale fermentation setups. These approaches not only enhance system performance but also deepen our understanding of how such machinery functions within the cellular environment. Additionally, these systems often incorporate other enzymes to improve adhesion to plastic surfaces, further boosting PET degradation efficiency. However, their reliance on inducers and antibiotics remains a limitation for optimal functionality.

In our study, we engineered an *E. coli* strain capable of displaying PETase and FAST-PETase on the outer membrane without the need for inducers or antibiotics, enabling constant transcription and translation of the target enzymes. This whole-cell biocatalyst exhibited activity against semi-crystalline PET powder, producing 0.18 mM of MHET at OD_600_=1, pH 7, over 3 days. Additionally, the *E. coli* MS92 strain retained up to 70% of its activity against BHET after 10 passages highlighting its stability and potential for sustainable PET degradation.

## Conclusions and outlook

Our approach, which integrates PET-degrading enzymes directly into the microbial genome, represents a significant advancement in developing self-contained, biosafe systems for plastic recycling. By harnessing Synthetic Biology and engineering enzymes with enhanced properties within modular whole-cell chassis, this approach provides a scalable and sustainable solution for large-scale, antibiotic-free fermentation processes. These engineered bacterial cell factories, with their genomic capability to convert PET plastic into valuable biomass or use it as a sole carbon source^44,45^, open new avenues for recycling plastics into useful products, paving the way for efficient and environmentally friendly plastic waste management.

## Materials and Methods

### Materials

PET powder (particle size < 300 μm, crystallinity >50%) was purchased from Goodfellow Co (Product Code ES30-PD-000132). BHET, MHET, and TPA was purchased from Sigma-Aldrich and were used as standard samples.

### Methods

#### Generation of Lpp deleted *E. coli* strain (MS33 strain)

The CRISPR-Cas9-assisted genome editing technique was employed to delete a 25 bp coding sequence from the endogenous Lpp gene while introducing a TAA stop codon, resulting in a truncated Lpp gene sequence within the genome of *E. coli* BL21 (DE3) cells, following a previously published method^30^. The pEcgRNA-ΔLpp(N20) plasmid was constructed by designing a 20 bp gRNA sequence that targets the Lpp coding region, with an NGG sequence at the 3’-ends, using the Golden Gate cloning method^46^. Initially, the pEcCas plasmid (Addgene #73227) was transformed into *E. coli* BL21 (DE3) cells, and the λ Red system genes were induced by adding 10 mM arabinose, 30 minutes before the cells reached an OD_600_ of 0.3-0.4 to prepare electrocompetent cells. Subsequently, 100 ng of pEcgRNA-Δlpp(N20) and 400 ng of donor DNA, containing 35 bp homology regions flanking the Cas9 cutting site and the premature stop codon, were co-electroporated into 100 µL of competent cells using a 1 mm gap electroporation cuvette, with a 2.5 kV pulse applied using an Eppendorf™ Eporator™. After a 1-hour recovery in SOB medium, the cells were plated on LB agar containing kanamycin and spectinomycin and incubated overnight at 37 °C.

Three randomly selected colonies were isolated and cultured overnight in LB medium with kanamycin. Colony PCR was performed to amplify the target Lpp region within the *E. coli* genome using primers complementary to sequences upstream and downstream of the target site. Sanger sequencing of the purified PCR products confirmed the accurate editing of the genomic region (Figure. S1). Finally, plasmid curing was conducted to remove the plasmids from the modified *E. coli* BL21 (DE3) cells, which are hereafter referred to as the *E. coli* MS33 strain.

#### Plasmid construction for genomic integration of Lpp-PETase

To construct the pDonor plasmids, the LPP promoter and signal sequence regions were initially amplified via PCR using the *E. coli* genome as the template. The vector backbone, containing the Right End (RE) and Left End (LE) sequences of the transposon, was amplified from the previously constructed pDonor plasmid (PM453). The plasmids pDo-ColA-Lpp-Cys-PETase-His (PM530), and pDo-Lpp-Cys-PETase-His-Cys-MHETase-c-Myc (PM532) were then constructed by amplifying the target enzyme gene fragments via PCR using Phusion™ High-Fidelity DNA Polymerase (Thermo Scientific™) and subsequently assembling them with the GeneArt™ Gibson Assembly HiFi Master Mix (Invitrogen™). The pDo-ColA-Lpp-Cys-FAST-PETase-His (PM570) was generated by making five site directed mutagenesis in wild-type PETase gene using PM530 as template. The authenticity of the constructed pDonor plasmids was confirmed through Sanger sequencing, with the construct maps provided in Supplementary Figure S8. All primers and oligonucleotides were synthesized by Integrated DNA Technologies (IDT).

#### Lpp-Cys-PETases mini transposon integration

A novel RNA-guided DNA integration tool mediated by CRISPR-associated transposases (Type I-F CASTs)^30^ was employed to insert Cys-PET-hydrolyzing gene cassettes, along with the LPP promoter—a strong constitutive promoter—into the *E. coli* genome at designated insertion sites specified in the crRNA array^33^. Initially, pTetEffector-array (PM452 or PM511)^31^ and pDonor plasmids (PM530, PM532, or PM570) were co-transformed into *E. coli* BL21 (DE3) chemically competent cells, which were then plated on LB agar containing kanamycin and streptomycin. The integration of the mini-transposon gene cassettes was achieved through the overexpression of transposition machinery proteins encoded by the pTetEffector-array plasmid. This process involved scraping the transformants and spreading appropriately diluted cells on freshly prepared LB agar plates supplemented with kanamycin, streptomycin, and 5.0 ng/mL anhydrotetracycline (aTc). After a 30-hour incubation at 37 °C, several colonies were randomly isolated and cultured overnight in LB liquid medium with kanamycin. Colony PCR was then performed to detect the insertion of the Mini-Tns (Lpp-Cys-PET-hydrolyzing gene cassettes) using pairs of primers flanking the target sites within the *E. coli* genome, as outlined in the crRNA array (Supplementary Figure S10). The expected PCR product sizes, confirmed through agarose gel electrophoresis, validated the successful integration of the target genes.

To remove plasmids from Mini Tn-integrated strains, the pCutamp plasmid (provided by Sheng Yang; Addgene plasmid #140632)^33^, which carries a gRNA sequence under the control of a rhamnose-inducible promoter that targets the AmpR promoter in the pTetEffector and pDonor plasmids, was chemically transformed into the cells. After a 1-hour recovery at 37°C, 0.3 mL of the recovered culture was transferred to 0.9 mL of LB medium containing 50 μg/mL ampicillin (Apr) and 10 mM L-rhamnose, then incubated at 37°C with shaking at 250 rpm for 5 hours. A 100 μL aliquot of this culture, diluted 10,000-fold, was spread on LB agar plates supplemented with 50 μg/mL ampicillin and 5 mM L-rhamnose, and incubated overnight. Individual colonies were selected for each strain and grown overnight in 0.5 mL of LB medium containing 10 mM sucrose. Subsequently, 100 μL of this culture, diluted 1,000,000-fold, was plated on LB agar with sucrose to isolate single colonies. The pCutamp plasmid encodes the SacB gene, which produces an enzyme that converts sucrose into levan, a substance highly toxic to *E. coli*. Plasmid elimination in the cured strains was confirmed by spotting individual colonies (3–5 per strain) onto LB agar plates containing kanamycin, spectinomycin, and ampicillin, as well as onto LB plates without antibiotics. Colonies that grew on LB plates without antibiotics but not on those with antibiotics were confirmed to have successfully undergone plasmid curing. The cured strains were subsequently grown in LB medium to produce glycerol stocks or competent cells.

#### Expression test of the PETase

The PETase expression was investigated using a plasmid-based system. *E. coli* Codon-optimized genes for PETase, and MHETase were synthesized in pUC57 universal cloning vector by GenScript and recloned into the pET29b expression plasmid, which is controlled by the T7 promoter and features a C-terminal His-tag. The plasmids were then transformed into *E. coli* BL21 (DE3) or *E. coli* MS33 cells. The following day, glycerol stocks of the transformed cells were prepared by combining 0.75 mL of the pre-culture with 0.75 mL of 50% sterile glycerol and storing the mixture at -80 °C.

For gene overexpression, 50 mL of LB medium in 250 mL Erlenmeyer flasks was inoculated with overnight cultures grown in LB+Kan medium to achieve an OD_600_ of 0.05. The flasks were incubated at 37 °C and 250 rpm for 2-3 hours until the cell density reached an OD_600_ of 0.6-1.0. Expression was then induced by adding 0.2 mM IPTG, and the cultures were incubated at various temperatures (15°C, 25°C, 37°C) to identify the optimal temperature for intracellular or outer-membrane-exposed PETase expression. Concurrently, cell samples were collected before and after induction for expression analysis using SDS-PAGE. After 20 hours of induction, the cells were harvested by centrifugation at 5,000 × g for 10 minutes at 4 °C.

For expressing PET-hydrolyzing genes in genetically modified cells, a glycerol stock was first inoculated into LB medium and cultured overnight at 37 °C. Then, 1% of this pre-culture was used to inoculate 50 mL of LB medium, which was incubated at 25 °C with shaking at 250 rpm for 20 hours or more. This approach allowed for evaluating how incubation time affects the expression of enzymes displayed on the cell surface. The cultured cells were harvested by centrifugation at 5,000 × g for 10 minutes at 4 °C for subsequent analysis.

#### SDS-PAGE and Western Blotting

To evaluate the expression of PET-hydrolyzing enzymes in *E. coli*, a 12% Tris-Glycine SDS-PAGE analysis was performed. Protein bands were visualized on the gels using Coomassie Brilliant Blue R-250 staining. Following electrophoresis, the protein bands were transferred from the SDS-PAGE gel onto a TransBlot® Turbo^TM^ Mini-size polyvinylidene fluoride (PVDF) membrane at a constant voltage of 100V for Western blotting. The PVDF membrane was pre-equilibrated in absolute ethanol for 2-3 minutes. After the transfer, the membrane was blocked with TBS buffer (Tris 20 mM, NaCl 150mM, pH 7) containing 5% skim milk powder for 30 minutes at room temperature (RT). Subsequently, GenScript THE™ His-Tag Monoclonal Antibody [HRP] was applied at a concentration of 60 ng/mL, and the membrane was incubated for 1 hour at RT. Finally, the membrane was washed three times with TBS buffer, and the targeted His-tagged enzymes were detected using the Immobilon® Forte Western HRP substrate solution. Proteins were visualized using the gel documentation system FluorChem Q (Cell Biosciences).

#### Membrane protein extraction

Membrane proteins were isolated using a modified version of the method outlined by Jarmander et al.^47^. Following enzyme expression, cells were collected by centrifugation at 5,000 × g for 10 minutes at 4°C to remove the LB medium. The resulting cell pellets were then washed and resuspended in buffer A (50 mM Tris-HCl, pH 7.5). Cell disruption was performed using a French® Pressure Cell Press SIM-AMINCO at 14,000 psi, with each sample passed through the instrument three times. To remove non-disrupted cells, the mixture was centrifuged at 3,500 × g for 10 minutes at 4°C. The supernatant was subsequently centrifuged at 30,000 × g for 40 minutes at 4°C to pellet the membrane-bound proteins. The supernatant, containing the soluble fraction, was stored at 4°C for later analysis, while the pellet, rich in membrane-bound proteins, was washed with buffer A. In the next step, the membrane-bound protein pellet was resuspended in buffer B (buffer A supplemented with 0.1% (v/v) sarcosyl N-laurylsarcosine sodium salt) and incubated at 4°C for 1 hour. The protein suspension was then subjected to centrifugation at 30,000 × g for 40 minutes at 4°C, allowing the separation of outer membrane proteins, which remained in the pellet, from inner membrane (IM) proteins in the supernatant. Finally, the outer membrane (OM) protein pellet was resuspended in buffer C (buffer A with the addition of 1% (v/v) Triton X-100 and 5 mM EDTA) and stored for subsequent analysis.

#### Flow cytometry analysis

Freshly harvested Cys-PETase-expressing cells were centrifuged at 1,700 × g for 10 minutes at 4°C. The resulting pellet was washed twice with PBS buffer. The cells were then resuspended in PBS buffer to achieve an OD_600_ of approximately 1.0. Next, a 100 µL aliquot of this cell suspension was incubated with GenScript THE™ His-Tag Antibody [FITC], mAb (10 µg/mL) for one hour at room temperature (RT), while protected from light with aluminum foil. After incubation, the sample was centrifuged again at 1,700 × g for 10 minutes at 4°C. The pellet was washed, resuspended in 200 µL of PBS buffer to reach an OD_600_ of 0.25, and stored at 4°C until analysis. The analyses were carried out in SH800S Cell Sorter, using a laser at 488 nm, a sorting chip, nozzle size 100 um, 100 000 event count.

#### Esterase activity assay against *p-*nitrophenyl alkanoates (*p*-NPAlk)

Enzyme activity was assessed using a UV-Visible spectrophotometer (Jasco J-770). An end-point assay was employed to evaluate the enzyme displayed on the cell surface. This evaluation utilized PETase’s ability to hydrolyze ester bonds in *p-*Nitrophenyl Alkanoates (*p-*NPAlk), including *p-*Nitrophenyl Acetate (*p-*NPAc, C2), *p-*Nitrophenyl Butyrate (*p-*NPB, C4), *p-* Nitrophenyl Octanoate (*p-*NPO, C8), and *p-*Nitrophenyl Dodecanoate (*p-*NPD, C12). The assays were performed in 500 µL reaction volumes, each containing 1 mM *p-*NPAlk (with a final acetonitrile concentration of 1%), a cell suspension with an OD_600_ of 0.04 in PBS buffer at pH 7.0 and 25°C. Reactions were initiated immediately upon the addition of the cell suspension. The reactions were halted after 5 min by centrifugation, and the supernatant was analyzed measuring the absorbance at 410 min nm (Abs_410_). To assess the linearity of enzyme activity, every 5 min, a 20 µL sample was taken, centrifuged for 1 minute, and 2 µL of the supernatant was analyzed using a Nanodrop at 410 nm. All samples and controls were analyzed in triplicate. The reactions and controls were run in triplicate and the concentrations of each sample were calculated using Beer-Lambert equation and applying ɛ= 7 800 M^-1^ cm^-1^ as extinction coefficient of *p*-nitrophenol^48^. One unit of *p*-NPAlk esterase activity was defined as the number of cells required to release 1 µmol of *p*-nitrophenol per minute under optimal conditions. The total cell concentration was obtained by serial dilution.

#### BHET and PET powder degradation measurement using HPLC

HPLC analysis was conducted using an XSelect CSM™ C18 5 µm column (4.5 × 100 mm, Waters Corp.) and a Photodiode-Array Detector (Waters Alliance e2695), which monitored the elution of BHET, MHET, and TPA at 240 nm, following the method outlined by Yoshida et al.^8^. The analysis was performed using an isocratic elution mode over a 20-minute period, with the mobile phase consisting of 80% 20 mM phosphate buffer (pH 2.5) and 20% methanol. After the elution, the column was washed with 100% methanol for 2 minutes and then re-equilibrated to the initial conditions for 1 minute. Each sample was injected in a volume of 50 µL. Sample preparation involved centrifugation at 13,000 × g for 10 minutes at room temperature. After centrifugation, 3.0 M HCl was added to adjust the pH to 2.5. To assess PET powder degradation by Cys-PETase expressed in *E. coli*, 5 mg of PET powder (Goodfellow Corp.) was incubated in 20 mM phosphate buffer (pH 7.0) with 1% DMSO at either 30°C or 37°C. Various cell concentrations (OD_600_: 0.5, 1.0, 3.0, and 6.0) were tested in a total reaction volume of 1 mL. At designated time points, the reactions were stopped by centrifugation. The supernatant was then collected, and the pH was adjusted to 2.5 using 3.0 M HCl. Standard curves of BHET, MHET, and TPA were obtained by HPLC using commercial standards to compare the retention times and quantify the concentrations (Figure S13-14).

#### Agar-plate activity assay

Following the protocol of Wang et al.^49^, with minor modifications, two types of plates were prepared to assess enzyme expression and localization. LB agar plates for plasmid-based expression tests were supplemented with 8 mM BHET, 0.2 mM IPTG, and 50 µg/mL kanamycin. In contrast, LB agar plates containing only 8 mM BHET were used to evaluate genetically modified cells. A 2 µL aliquot from overnight cultures of *E. coli* BL21(Cys-PETase) and *E. coli* MS33(Cys-PETase) strains, as well as *E. coli* MS75 and *E. coli* MS92 strains, was plated. The plates were then incubated at 25°C for 3 days to assess BHET degradation and halo formation.

#### Expression stability analysis

To compare the stability of enzyme expression between the pET29b-Lpp-PETase plasmid system and genomically integrated cells, we maintained consistent expression conditions as described previously. Cells from the initial passage were labeled as the P1 sample. Cultures were then initiated from these G1 samples and propagated through 10 generations (G1-G10). After each generation, cells were harvested, washed, and stored in a mixture of PBS and 25% glycerol at - 80°C until the activity assays were conducted.

For the enzyme assays, samples stored at -80°C were gently thawed and washed twice with 20 mM phosphate buffer (pH 7.0). The washed cells were centrifuged at 5,000 x g for 5 minutes at 4°C and resuspended in 20 mM phosphate buffer to achieve the desired OD_600_. Reactions were performed in 1 mL volumes using 1 mM *p-*NPAc (diluted in 1% acetonitrile) or 1 mM BHET (diluted in 1% DMSO) and incubated at room temperature for specified time intervals. The reactions were stopped by centrifugation. All assays, including controls, were conducted in triplicate.

#### Next-Generation Whole Genome Sequencing

The *E. coli* MS75 strain (BL21::Lpp-Cys-PETase-His) was selected for next-generation whole genome sequencing, conducted by SeqCoast Genomics LLC. After the strain was cultured in LB medium, genomic DNA was extracted using the PureLink™ Microbiome DNA Purification Kit (ThermoFisher Scientific). Short-read whole genome sequencing was then performed on an Illumina platform, generating 400 Mbp of data from 2.7 million reads, each 150 bp in paired-end format. To identify indels and point mutations, the Breseq computational pipeline was used to compare the *E. coli* MS75 genome against reference sequences from the *E. coli* BL21 (DE3) chromosome (CP053602) and the PM530 plasmid. Breseq provides detailed reports in annotated HTML format, highlighting single-nucleotide mutations, point insertions and deletions, large deletions, and new junctions caused by transposons. The analysis was conducted on the Digital Research Alliance of Canada server, with a summary of the results presented in Figures S15.

### Statistical analyses

Enzyme kinetics analyses employing chromogenic substrates and HPLC measurements were conducted on at least three independent sets of triplicate samples. The resulting data were then analyzed statistically using GraphPad Prism 10.1.2 software.

## Acknowledgments

Funding support for this work was provided by a National Research Council (NSERC) Discovery grant (RGPIN-2023-04787) held by Dr. David Levin. Dr. Nediljko Budisa and Dr. Hamid Reza Karbalaei-Heidari thank the Canada Research Chairs Program (Grant Nr. 950-231971) and Dr. Katherine Romero-Orejon thanks the Natural Sciences and Engineering Research Council (NSERC) of Canada through the Discovery Grants (RGPIN-04945-2017 and RGPIN-05669-2020) for support. Special thanks to Dr. Gerstein from the Department of Microbiology for the facility with the Flow cytometry analysis and Dr. McKenna from the Department of Chemistry for their assistance with Western Blot. We thank the members of Dr. Budisa Group and Dr. Levin Group for their assistance.

