## Supplementary Information for "Antibiotic-free whole-cell biocatalytic fermentation: *Escherichia coli* with surface-displayed PETases for sustainable plastic degradation"

[**Figure S8.** **a**) Map of pDonor plasmid PM530 (Lpp-Cys-PETase transposon) designed for genomic integration into *E. coli* BL21 (DE3) strain. The plasmid carries a ColA origin of replication to maintain a low copy number during transposition, thereby reducing experimental toxicity. The construct includes the Lpp promoter, and a 21-amino acid Lpp signal peptide positioned upstream of the PETase coding sequence, forming a DNA cassette flanked by RE and LE transposon sequences. This configuration enables constitutive expression of membrane-exposed PETase. **b**) In the PM532 plasmid, the Lpp signal sequence and codon-optimized MHETase coding sequence, with a C-Myc tag, were added downstream of PETase in a polycistronic format. The PM570 plasmid was constructed by introducing five mutations into the PETase coding sequence in PM530, creating the FAST-PETase donor plasmid.	11](#_Toc183031153)

[**Figure S10.** PCR Screening of **a**) *E. coli* MS75 and **b**) *E. coli* MS92 strains Following Transposition Experiment. In these experiments, the pTetEffector-array1 plasmid (PM452), containing 9 crRNAs, was used to induce transposition of the donor plasmid PM530 (pDo-ColA-Lpp-Cys-PETase-His) into the *E. coli* BL21 (DE3) genome, targeting the *E. coli* MS75 strain. The pEffector-array2 plasmid (PM511), containing 10 crRNAs, facilitated the transposition of PM570 into the genome of *E. coli* BL21 (DE3) for the *E. coli* MS92 strain. Successful integration of the Lpp-Cys-PETase transposon is indicated by a 2312 bp PCR product at site #7 in *E. coli* MS75 (downstream of the kdgA gene) and at site #10 in *E. coli* MS92 (downstream of a hypothetical gene, ECD_03539). Non-specific PCR products were detected at site #4 in crRNA-array1 and at sites #3 and #4 in crRNA-array2.	13](#_Toc183031155)

**a**

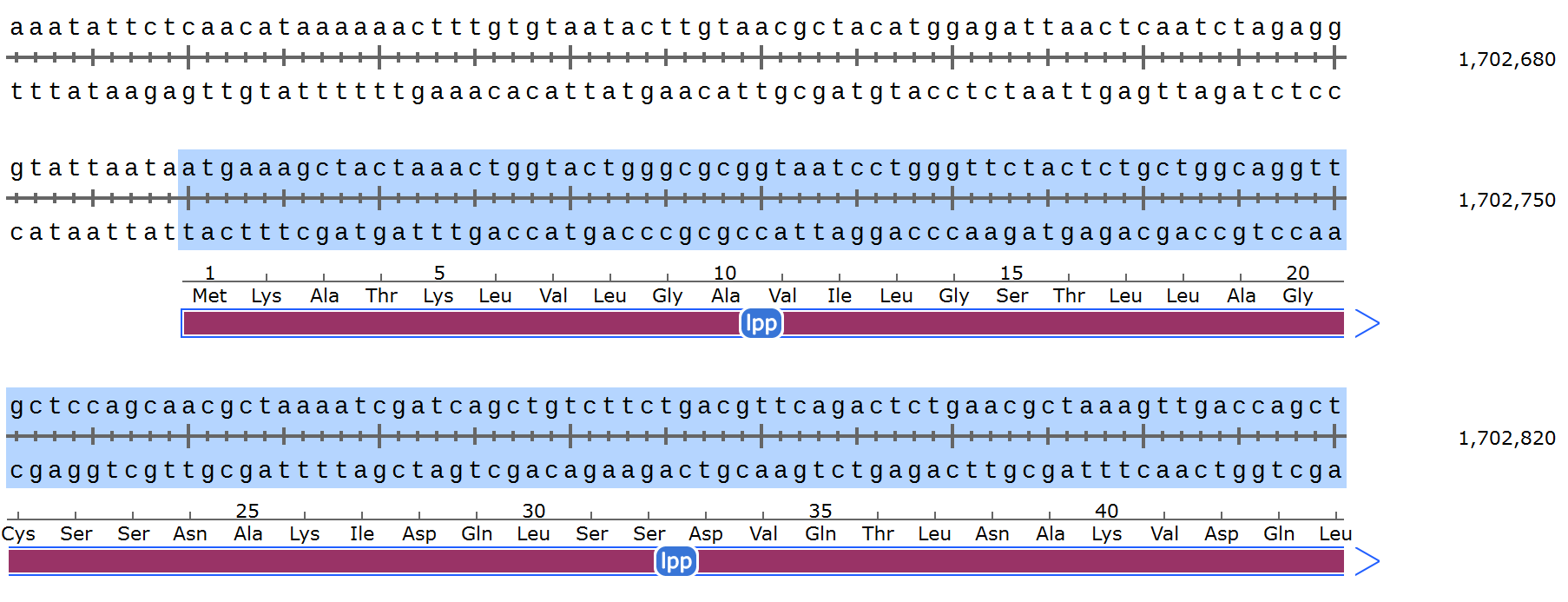

*E. coli* BL21 (DE3) (CP001509) Genome Sequence

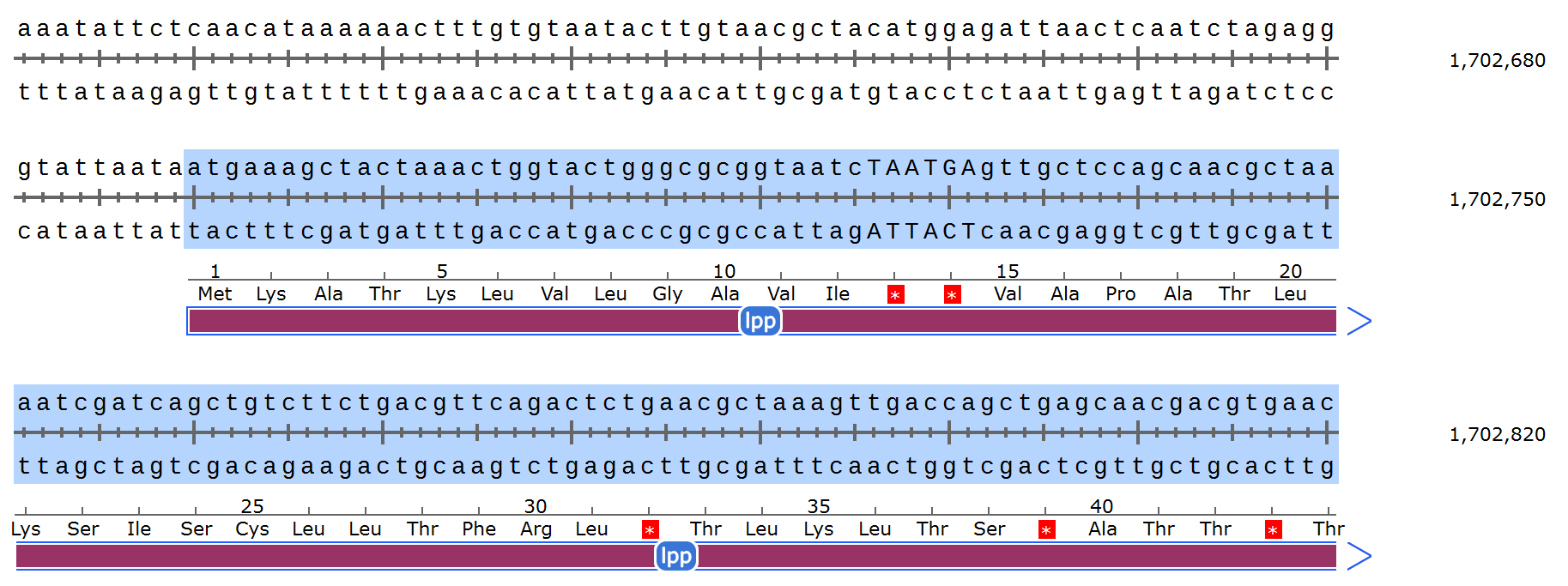

*E. coli* MS33 strain (Pre-mature Lpp) Genome Sequence

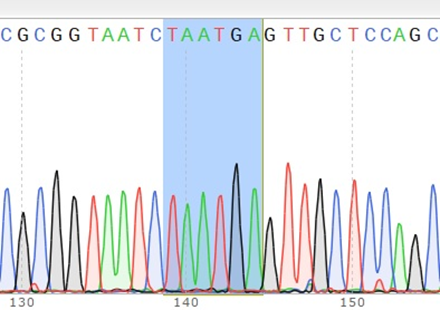

**b**

**Figure S1.** The Lpp gene sequence from *E. coli* BL21 (DE3). **a**) (Accession number: CP001509) before (wild-type) and after CRISPR-Cas9-assisted genome editing. **b**) Chromatogram of the target region after sequencing the PCR product from the Lpp region in the *E. coli* MS33 strain genome.

**b**

**c**

**a**

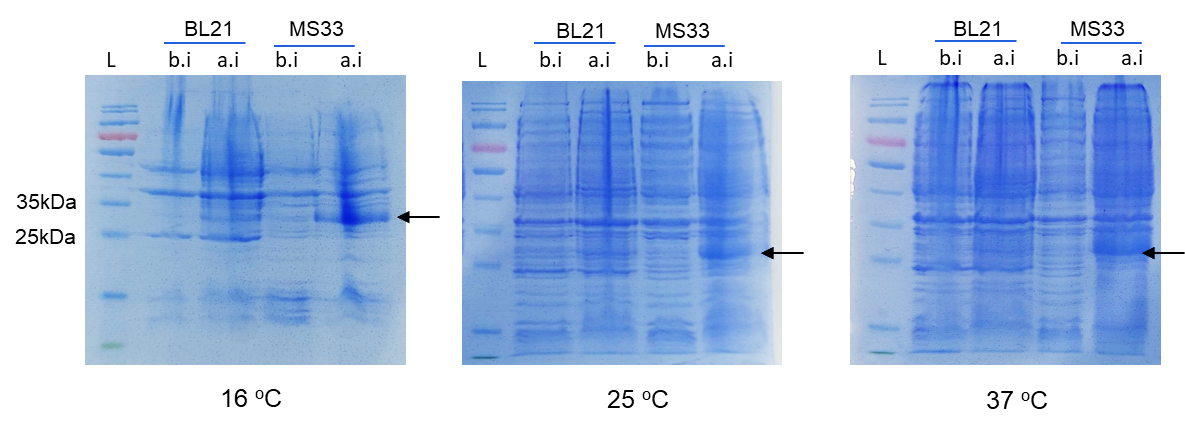

**Figure S2.** SDS-PAGE analysis of the effect of induced temperature on of Cys-PETase expression. The induction was performed at **a**) 16 ^o^C, **b**) 25 ^o^C and **c**) 37 ^o^C. L: ladder, b.i.: before induction, a.i.: after induction.

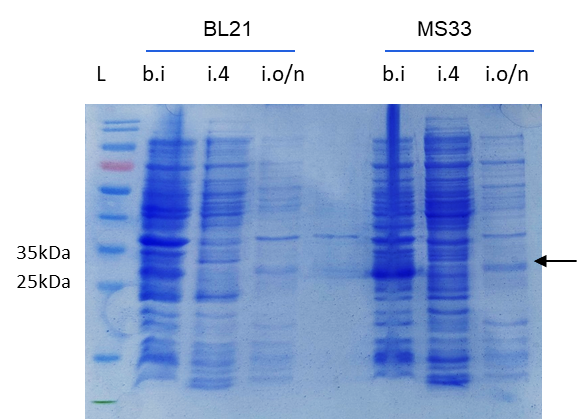

**Figure S3.** SDS-PAGE analysis of the effect of the time of induction of Cys-PETase expression at 25 °C. L: ladder, b.i.: before induction, i.4.: induction for 4h, i.o/n: induction of overnight.

|  | 1 d | 2 d | 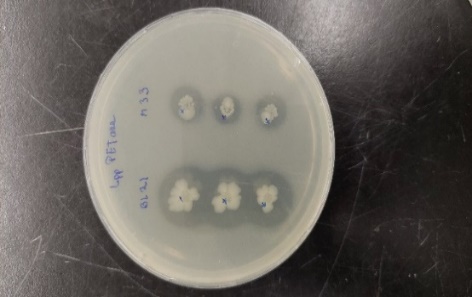3 d |
| --- | --- | --- | --- |
| **a** Cys-PETase | 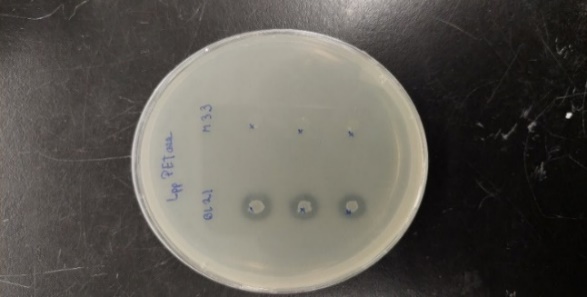 | 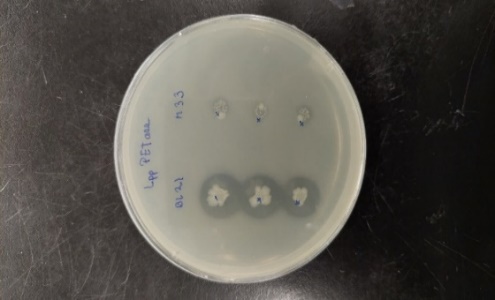 |  |
| **b** PETase | 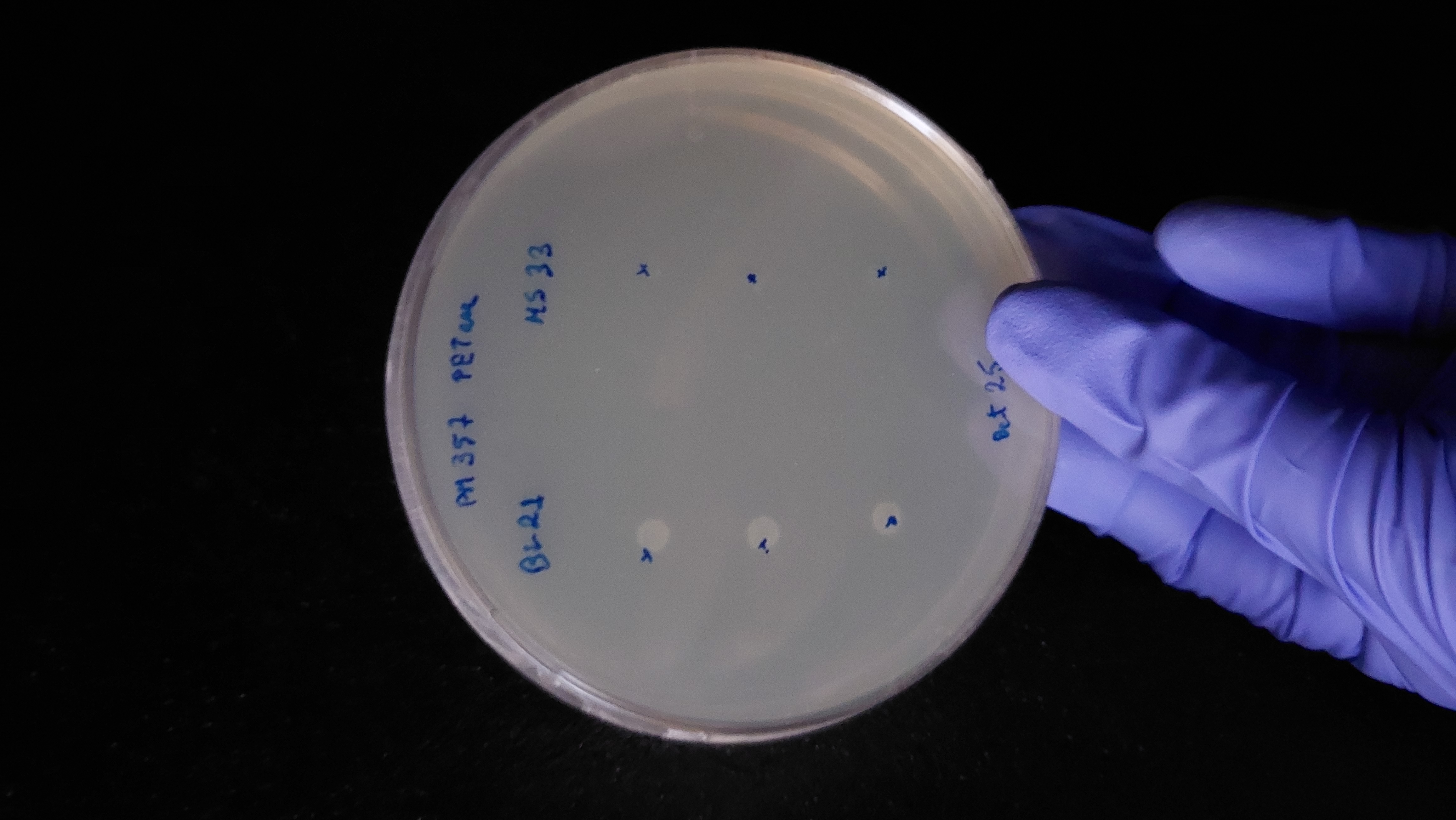 | 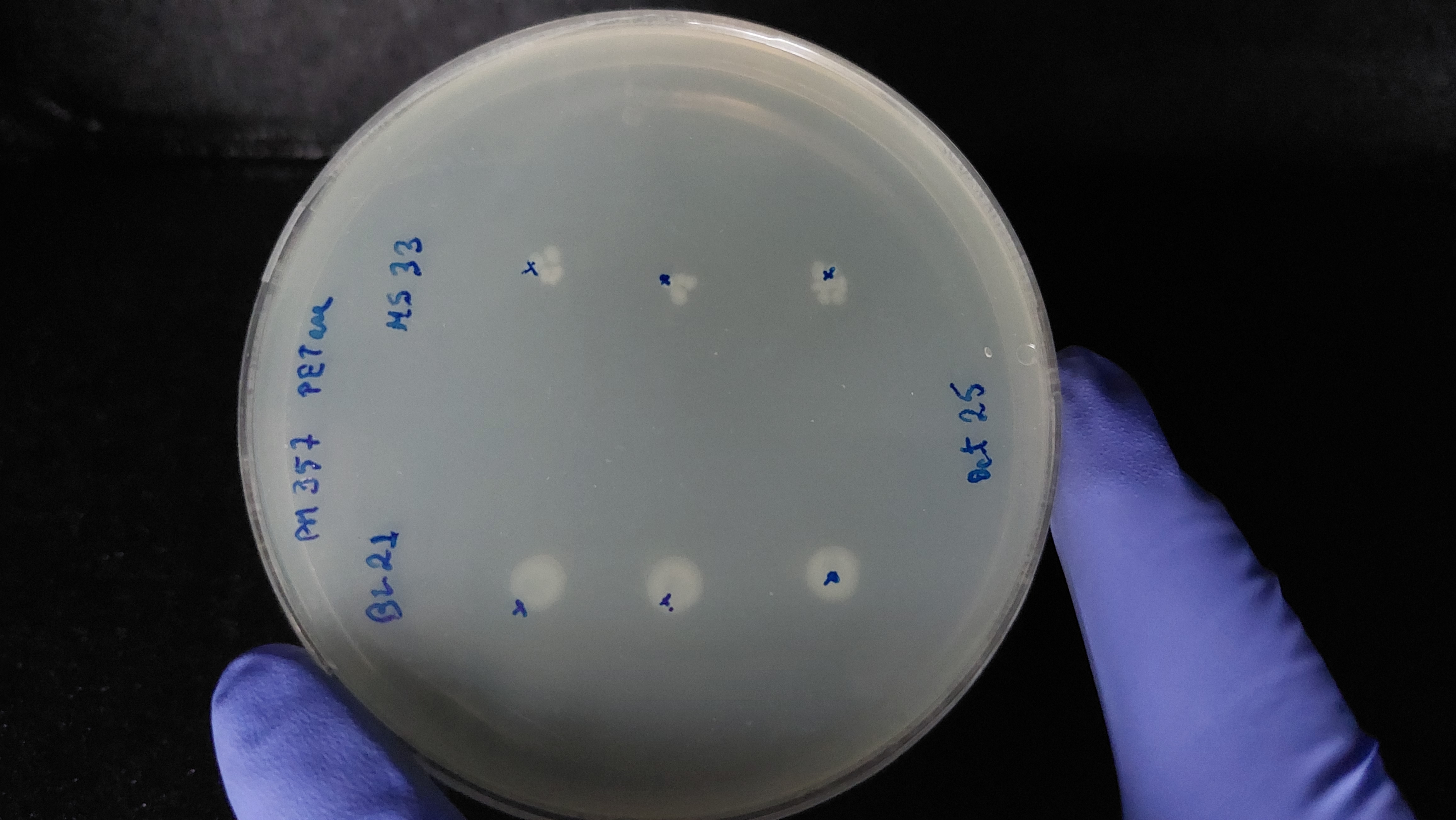 | 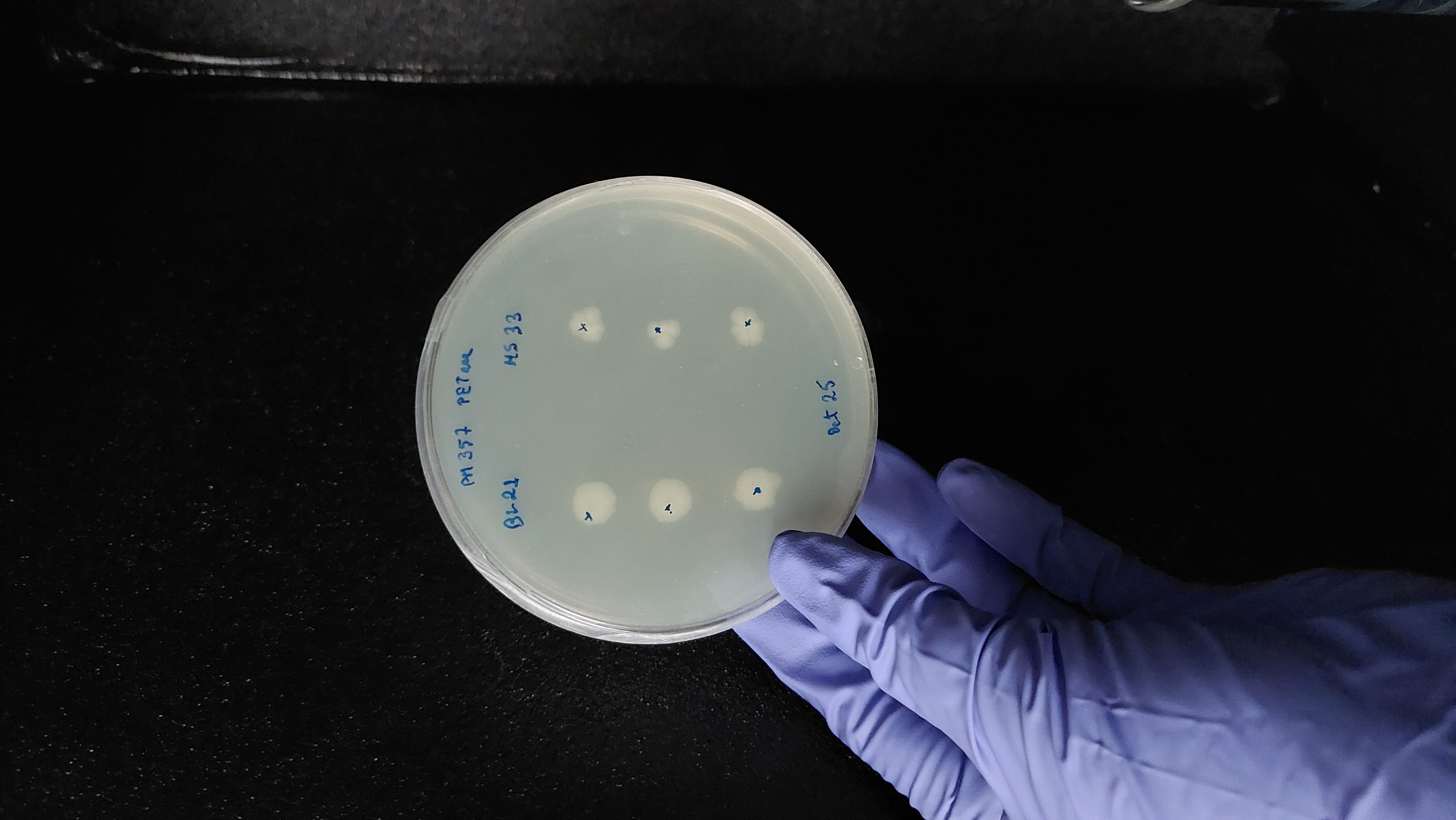 |
| **c** No plasmid | 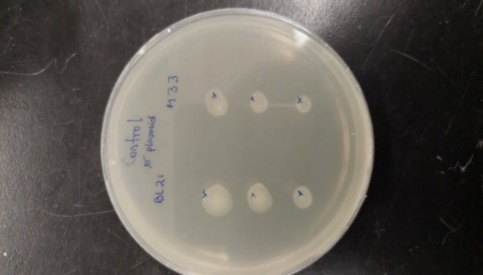 | 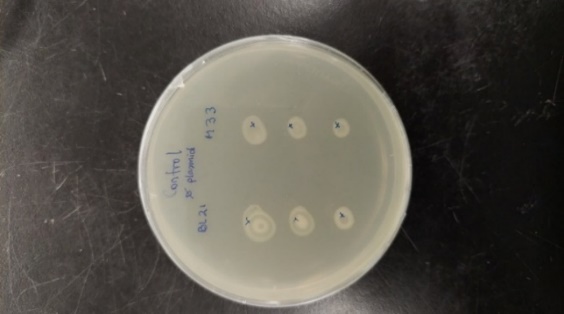 | 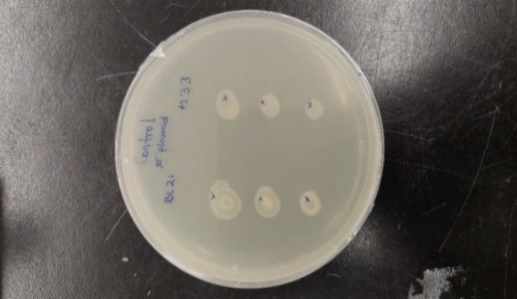 |

**Figure S4.** Comparison of halo formation in BHET-LB agar plates with different strains over 3 days of incubation. Samples were inoculated in triplicate. **a**) *E. coli* BL21 (Cys-PETase) and *E. coli* MS33 (Cys-PETase), **b**) *E. coli* BL21 (PETase) and *E. coli* MS33 (PETase) and **c**) *E. coli* BL21 and *E. coli* MS33 without plasmid.

|  | 1d | 7d | 13d |
| --- | --- | --- | --- |
| **a** PETase | 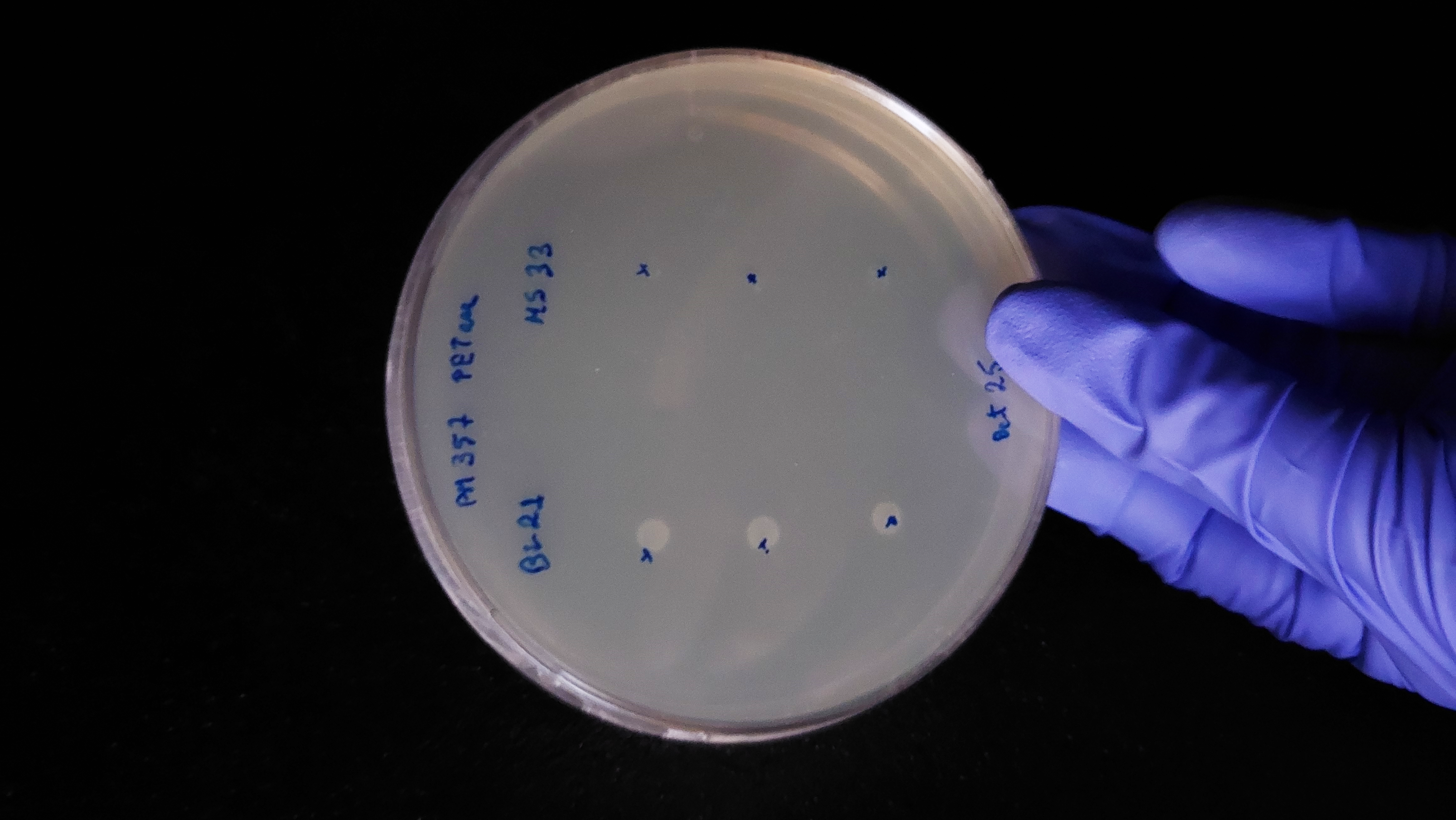 | 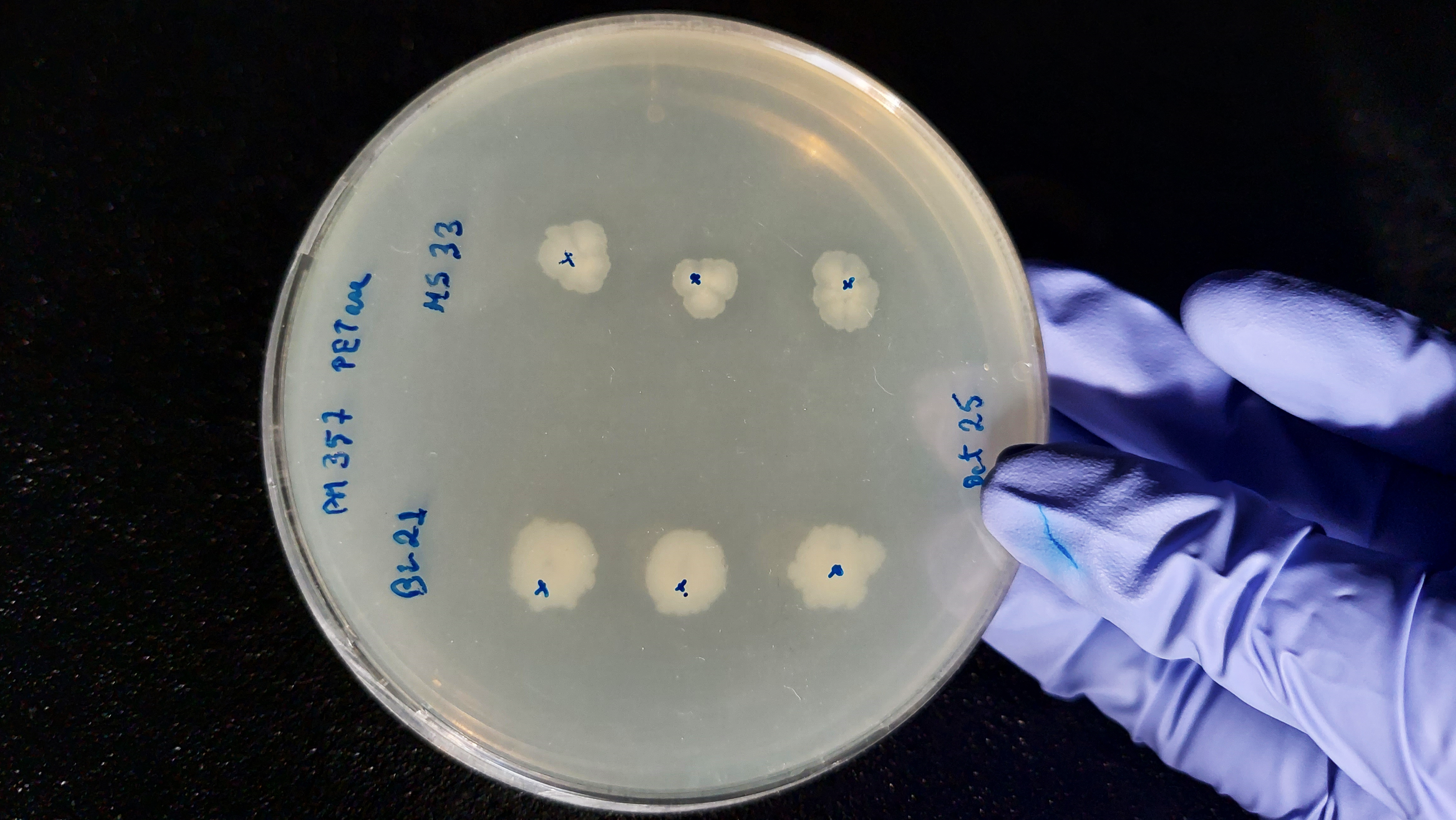 | 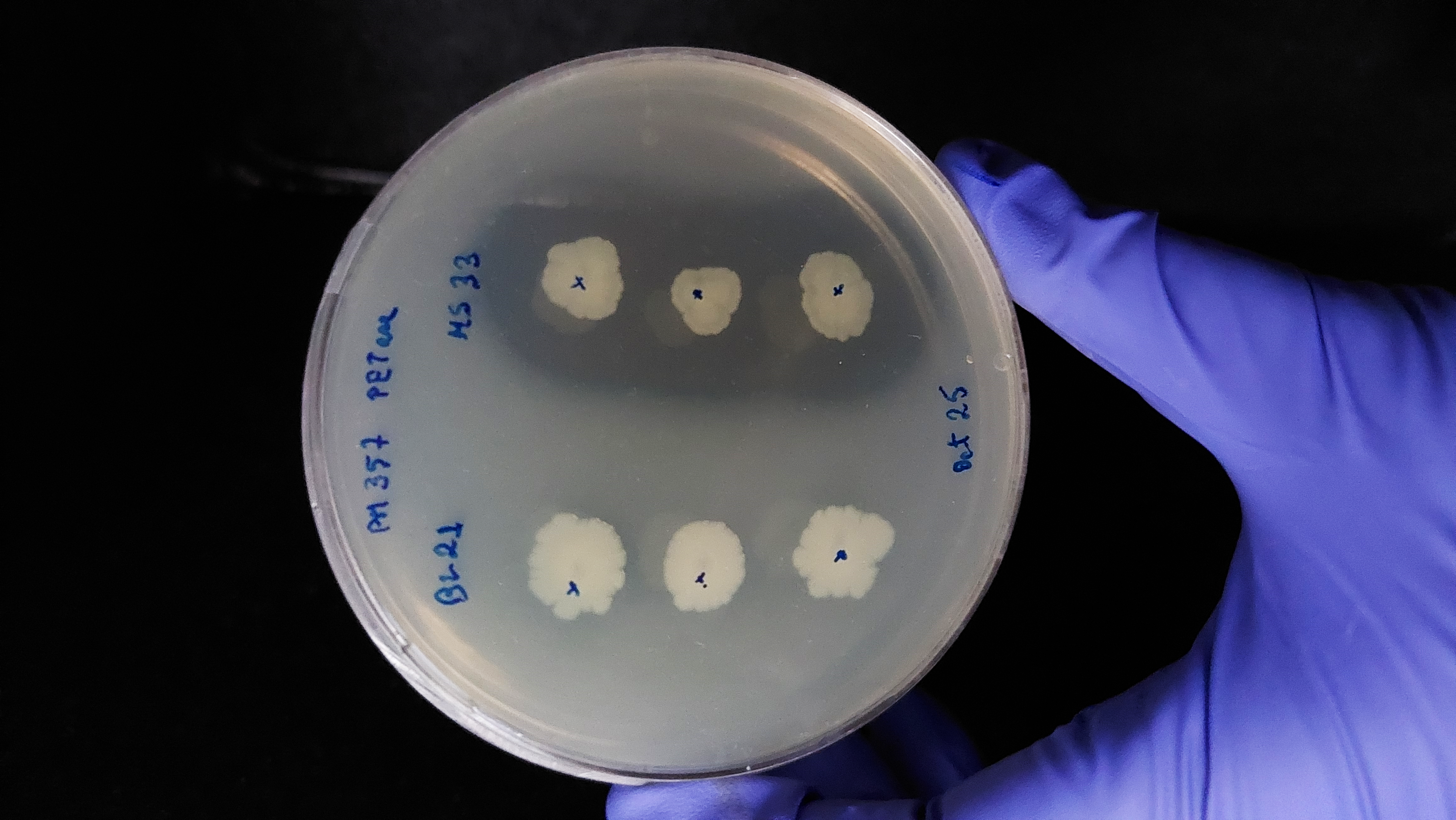 |
| **b** No plasmid | 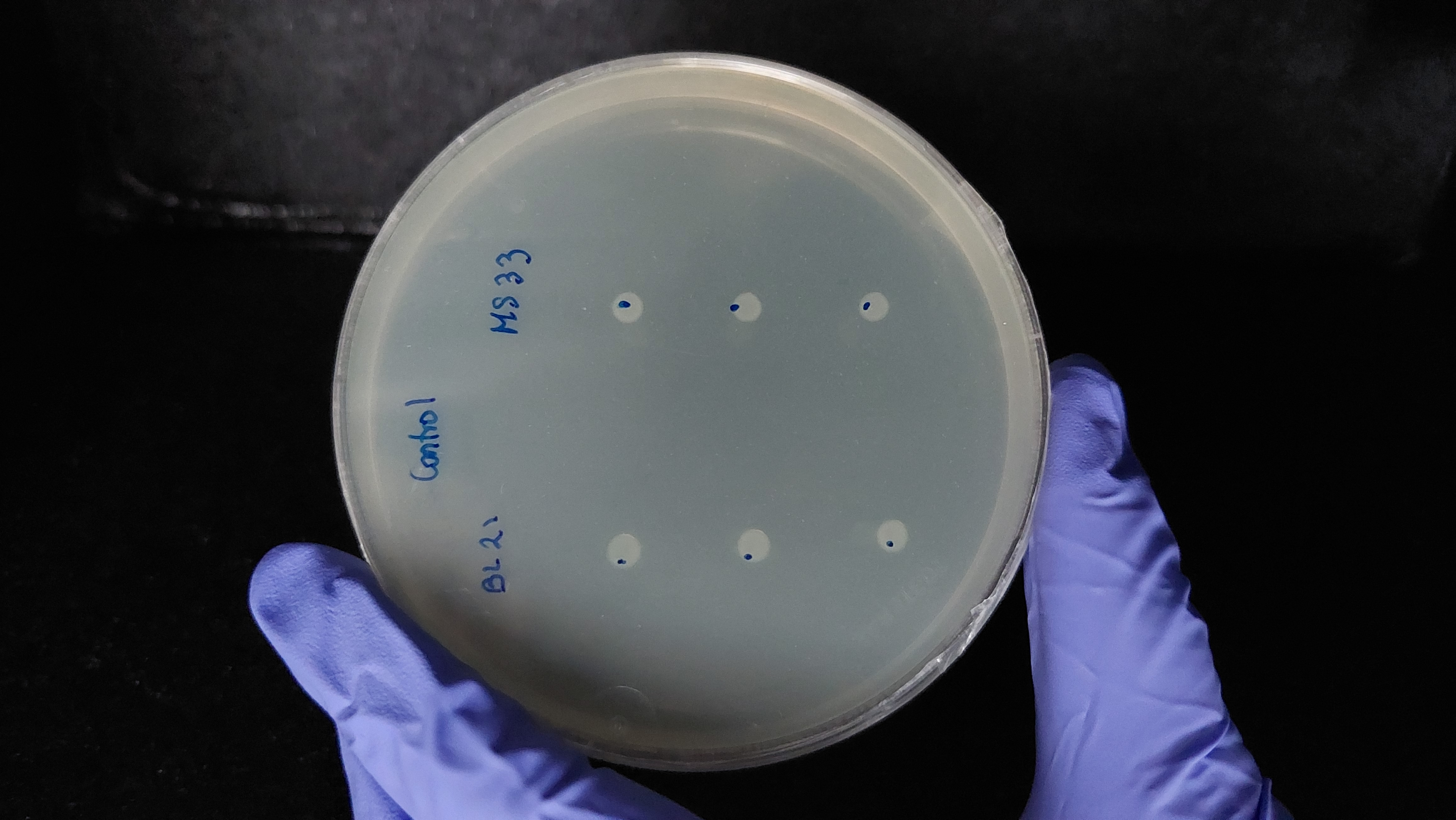 | 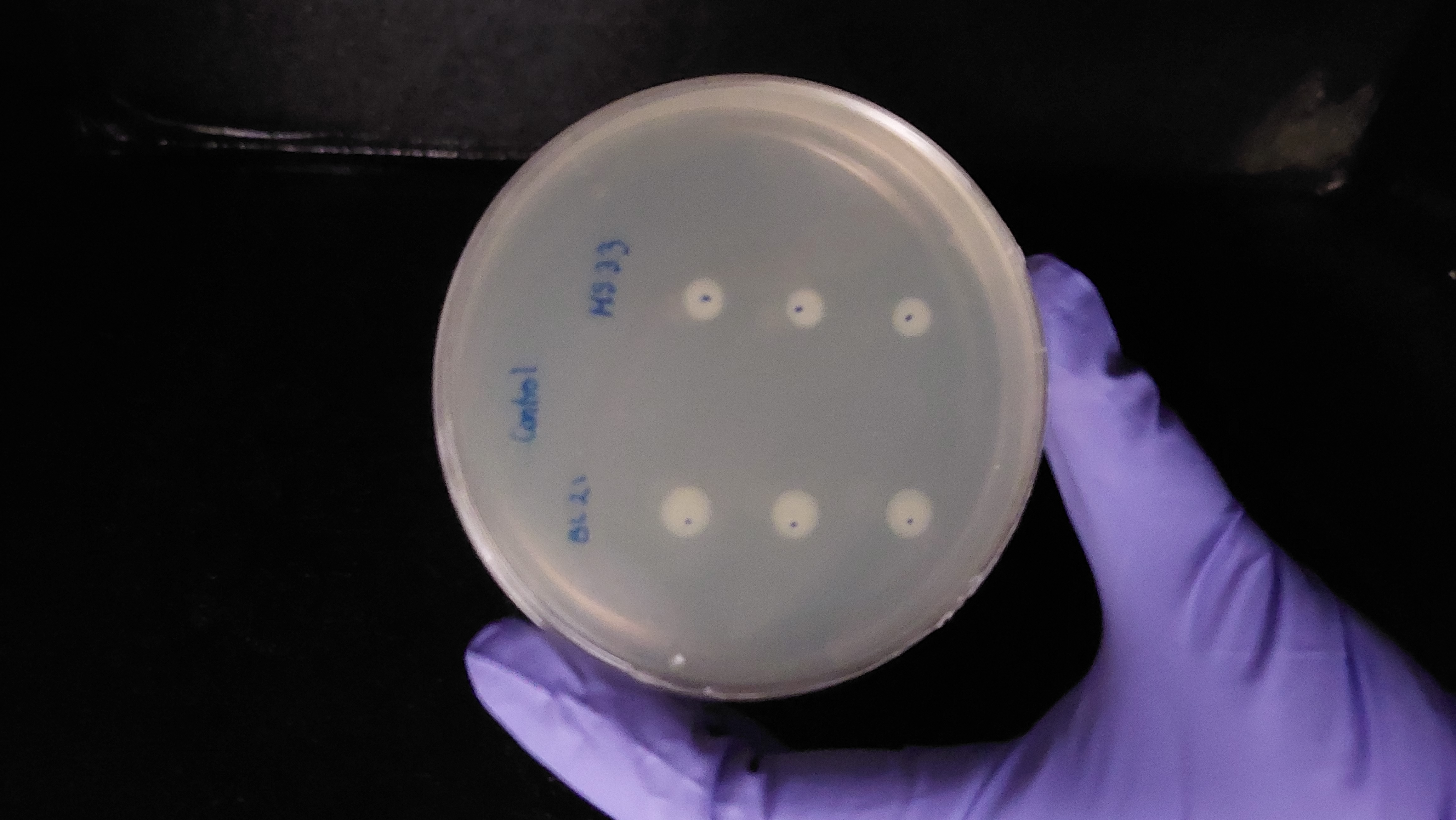 | 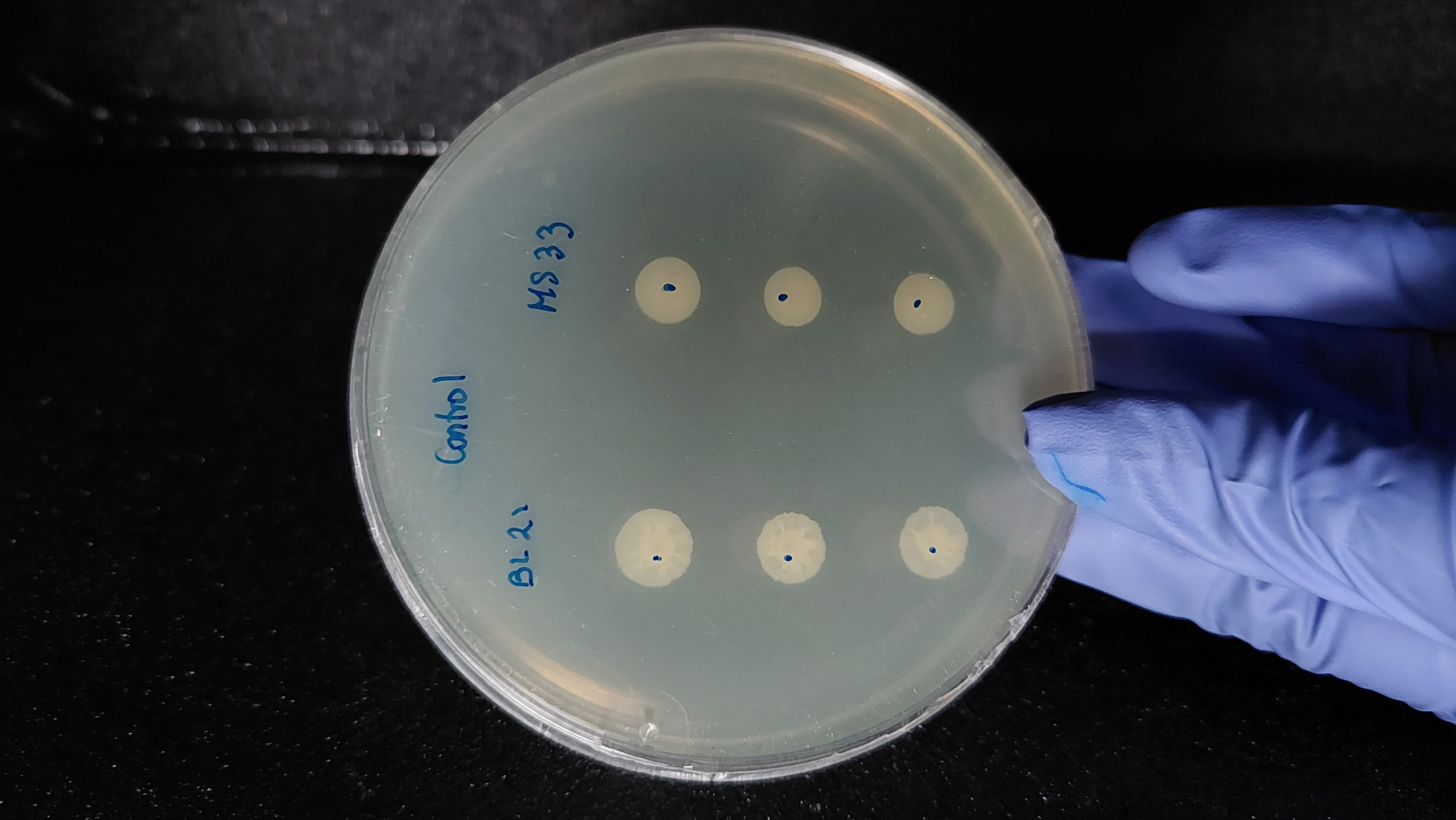 |

**Figure S5.** Comparison of the membrane permeability of different strains was assessed by observing halo formation on BHET-LB agar plates. Samples were inoculated in triplicate and incubated for 13 days. **a**) *E. coli* BL21 (PETase) and *E. coli* MS33 (PETase), and **b**) *E. coli* BL21 and E. *coli* MS33 without plasmid.

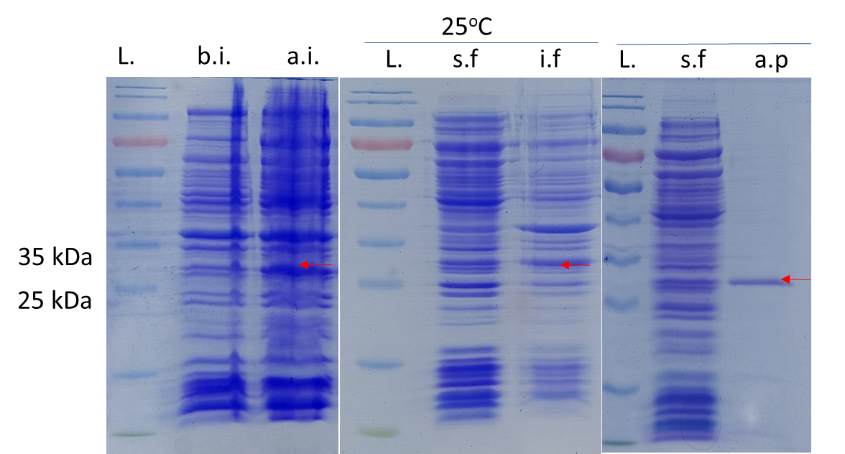

**Figure S6.** SDS-PAGE analysis of *E. coli* MS33(PETase) expression at 25 ^o^C. L: ladder, b.i.: before induction, a.i.: after induction, s.f.: soluble fraction, i.f.: insoluble fraction, a.p.: after Ni-column purification.

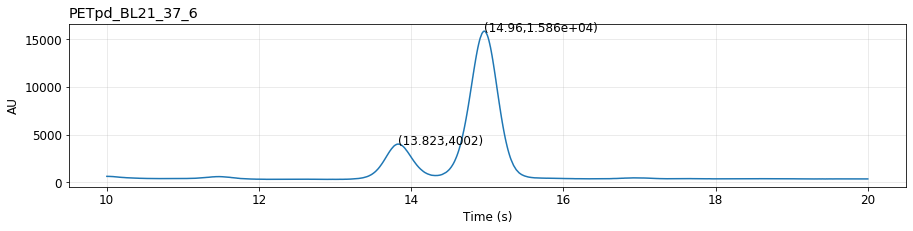

**Figure S7.** Chromatogram obtained by High-performance liquid chromatography-photodiode array (PDA) detector (HPLC-PDA). The peak at 14.96 min showed the presence of MHET as a product after incubation of *E. coli* BL21(Cys-PETase), OD_600_: 6, against PET powder at 37 °C for 24h.

**a**

**b**

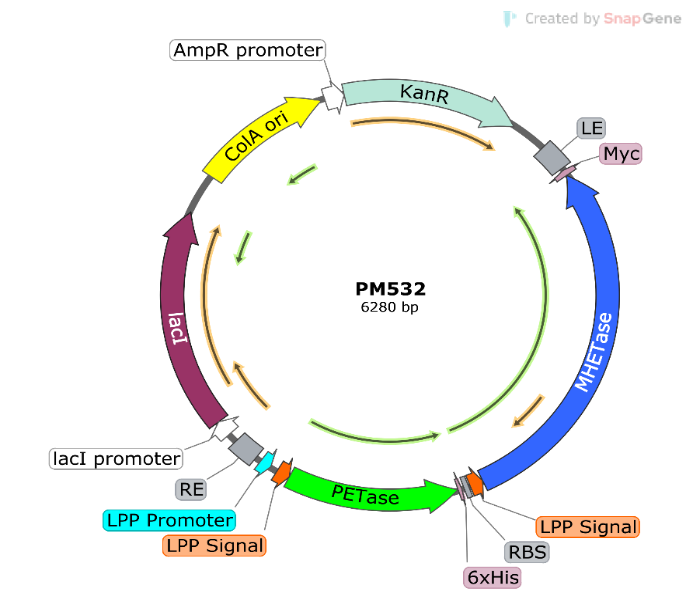

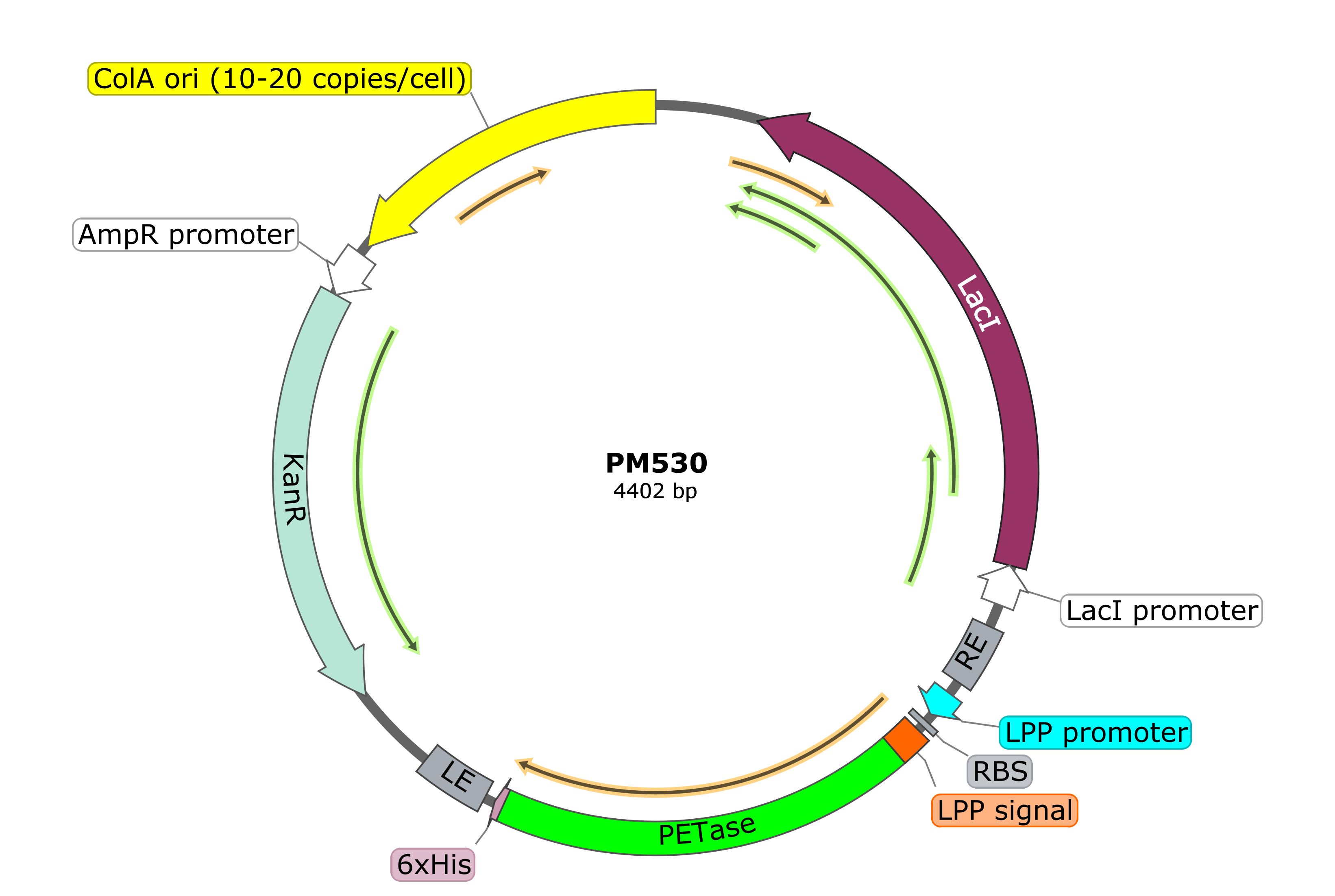

**Figure S8.** **a**) Map of pDonor plasmid PM530 (Lpp-Cys-PETase transposon) designed for genomic integration into *E. coli* BL21 (DE3) strain. The plasmid carries a ColA origin of replication to maintain a low copy number during transposition, thereby reducing experimental toxicity. The construct includes the Lpp promoter, and a 21-amino acid Lpp signal peptide positioned upstream of the PETase coding sequence, forming a DNA cassette flanked by RE and LE transposon sequences. This configuration enables constitutive expression of membrane-exposed PETase. **b**) In the PM532 plasmid, the Lpp signal sequence and codon-optimized MHETase coding sequence, with a C-Myc tag, were added downstream of PETase in a polycistronic format. The PM570 plasmid was constructed by introducing five mutations into the PETase coding sequence in PM530, creating the FAST-PETase donor plasmid.

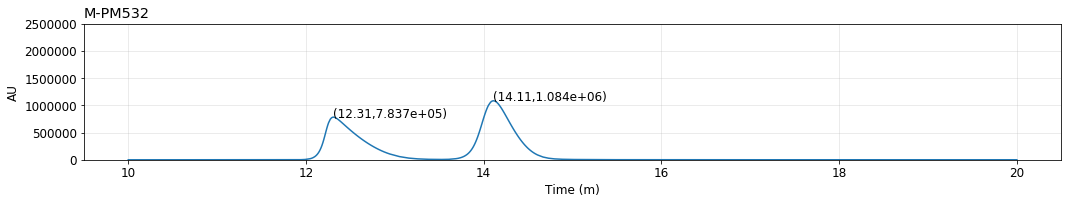

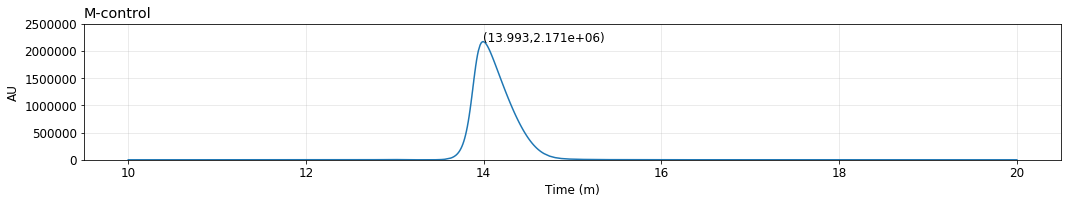

MHET

MHET

TPA

**Figure S9.** Chromatograms obtained by High-performance liquid chromatography-photodiode array (PDA) detector (HPLC-PDA). (**Top**) Peak at 13.99 min showing MHET as a negative control. (**Below**) Peak at 14.11 min shows the remaining amount of MHET after incubation of with *E. coli* BL21(PM532) cell harboring pDo-Lpp-Cys-PETase-Cys-MHETase (OD_600_: 1) at 37 °C for 1h and TPA product at 12.31 min.

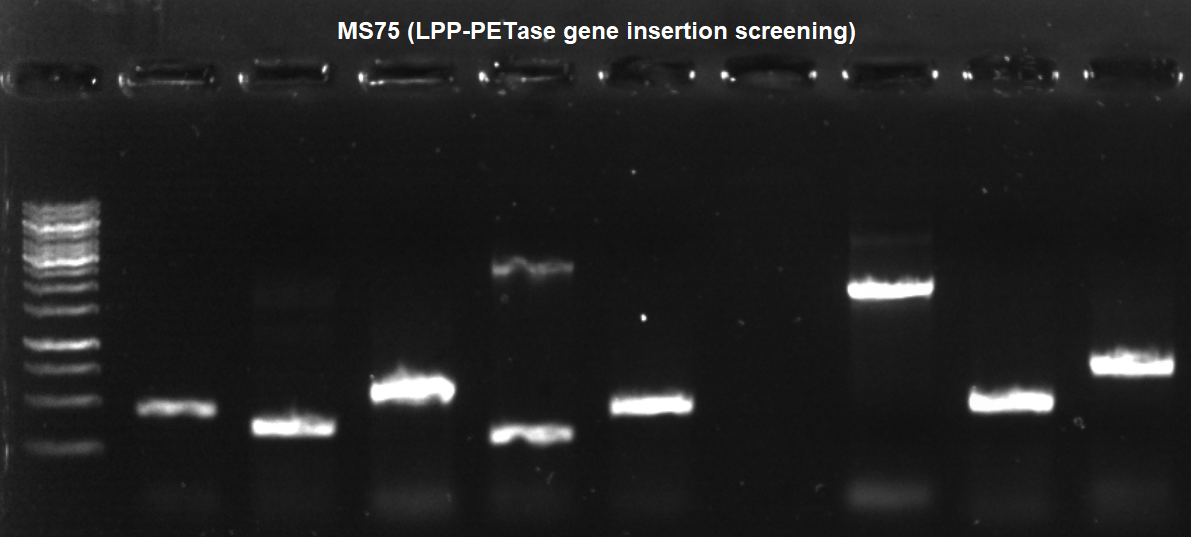

**a**

**b**

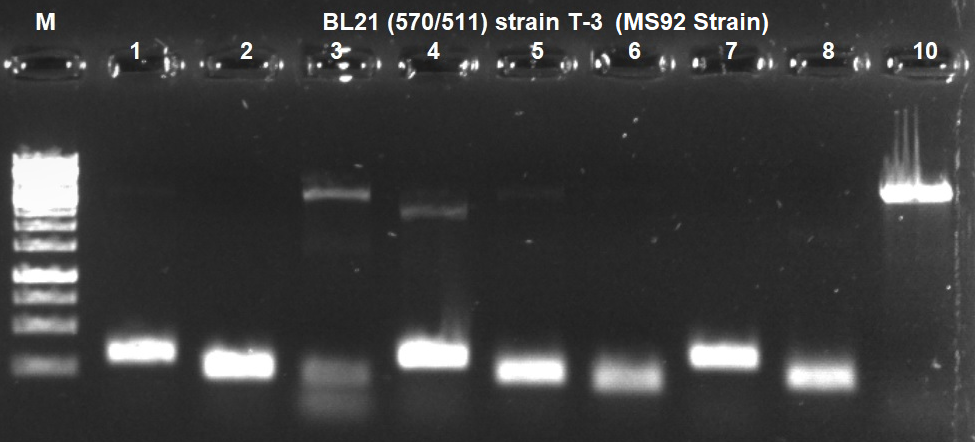

**Figure S10.** PCR Screening of **a**) *E. coli* MS75 and **b**) *E. coli* MS92 strains Following Transposition Experiment. In these experiments, the pTetEffector-array1 plasmid (PM452), containing 9 crRNAs, was used to induce transposition of the donor plasmid PM530 (pDo-ColA-Lpp-Cys-PETase-His) into the *E. coli* BL21 (DE3) genome, targeting the *E. coli* MS75 strain. The pEffector-array2 plasmid (PM511), containing 10 crRNAs, facilitated the transposition of PM570 into the genome of *E. coli* BL21 (DE3) for the *E. coli* MS92 strain. Successful integration of the Lpp-Cys-PETase transposon is indicated by a 2312 bp PCR product at site #7 in *E. coli* MS75 (downstream of the kdgA gene) and at site #10 in *E. coli* MS92 (downstream of a hypothetical gene, ECD_03539). Non-specific PCR products were detected at site #4 in crRNA-array1 and at sites #3 and #4 in crRNA-array2.

|  | Relative activity (%) |
| --- | --- |
| NC | 9.87279 |
| BL21(PETase) | 24.7369 |
| MS75-1d | 56.8249 |
| MS75-2d | 67.9149 |
| MS75-3d | 68.5375 |
| BL21(Cys-PETase) | 99.8436 |

**Figure S11.** Comparison of the esterase activity of the *E. coli* MS75 strain during 3 days of growth in LB medium as well as *E. coli* BL21 cells harboring the plasmid pET29b with the corresponding enzyme.

| 1 d | 3 d | 7 d | 13 d |
| --- | --- | --- | --- |
| 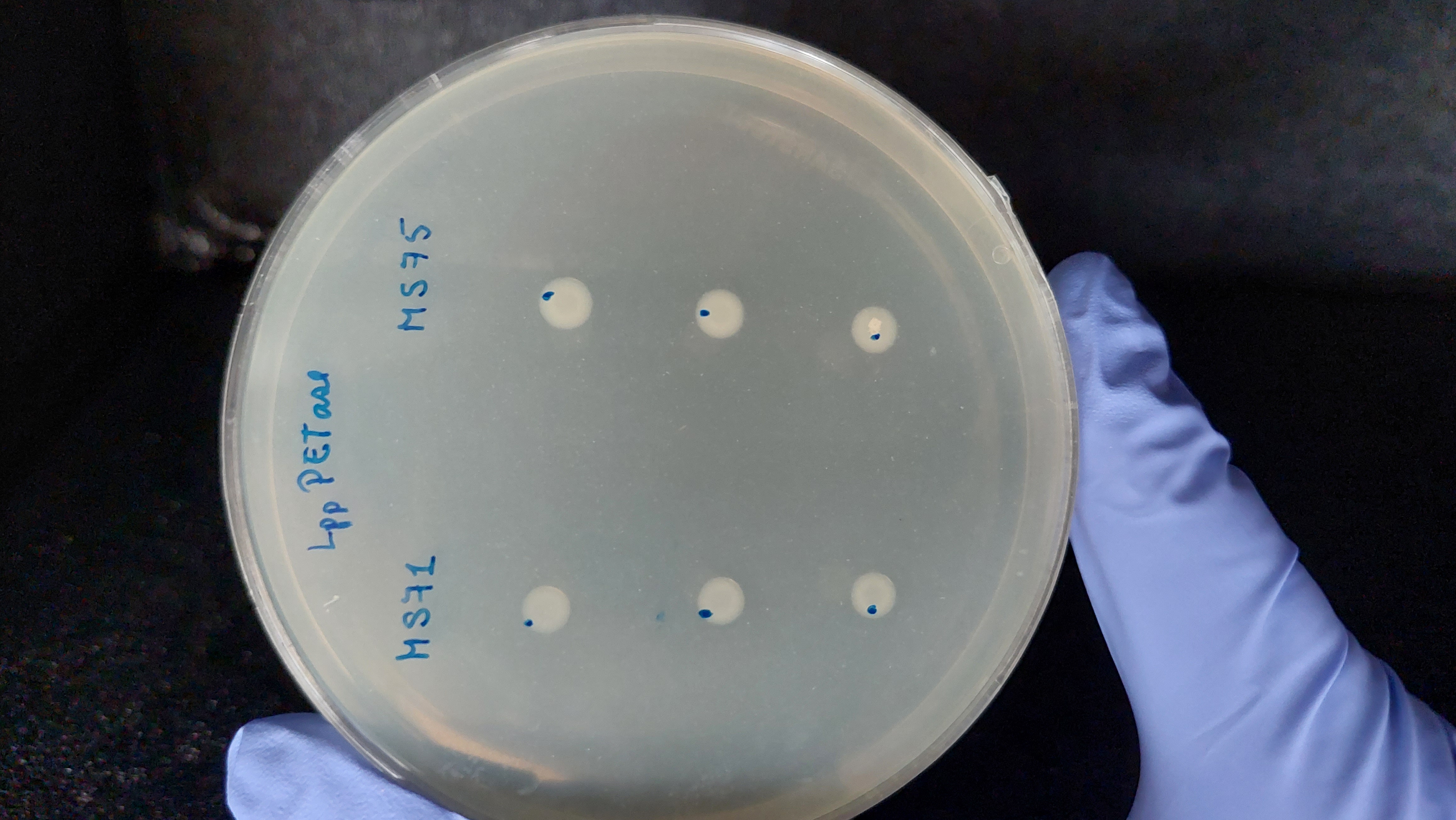 | 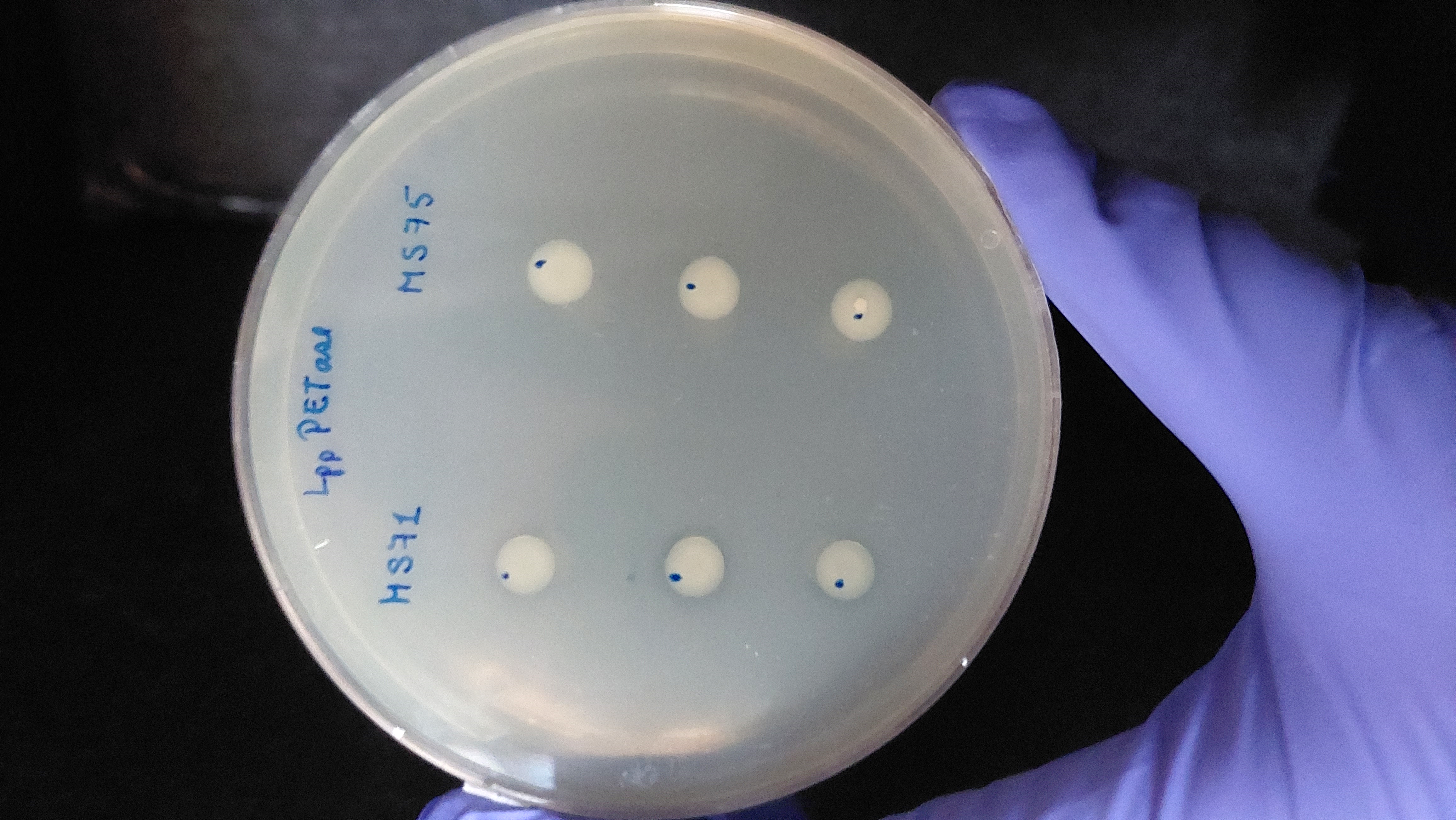 | 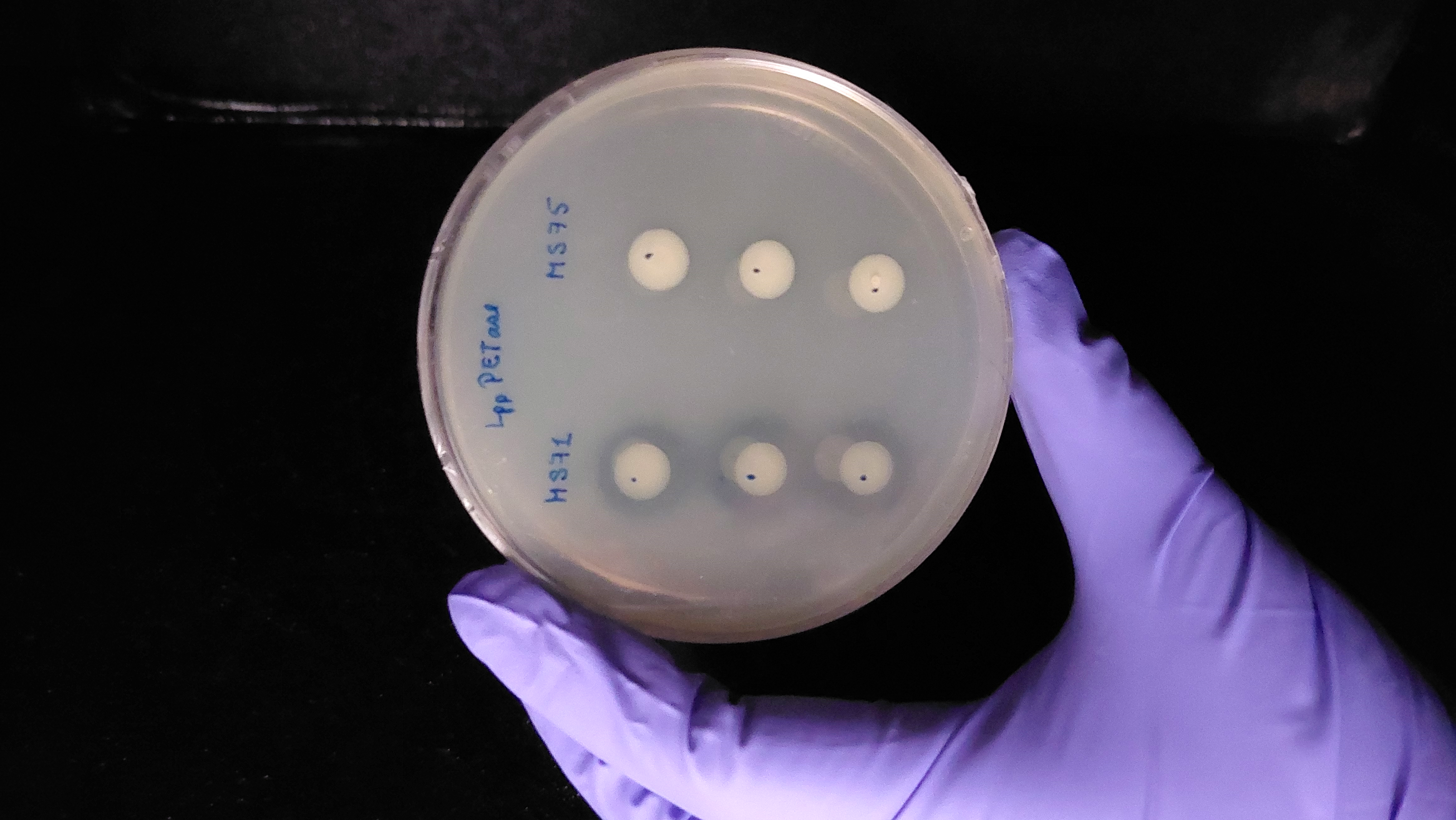 | 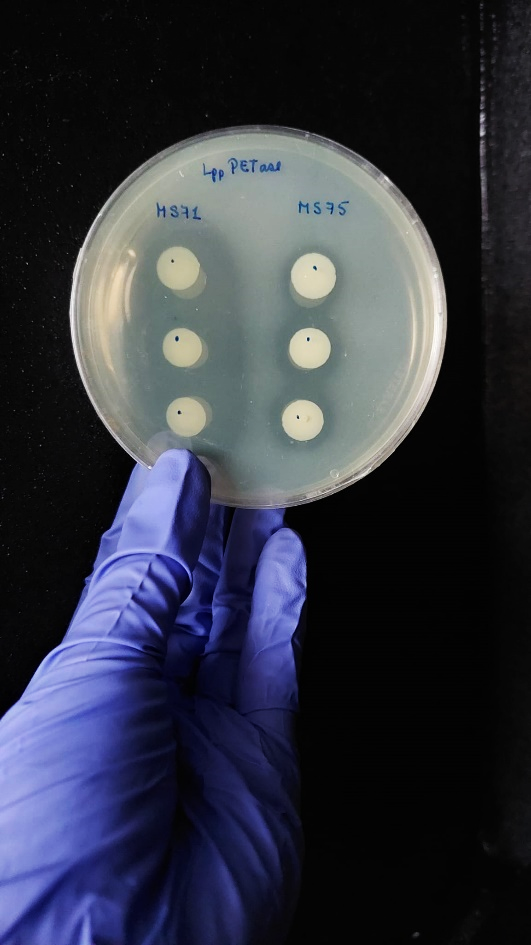 |

**Figure S12.** BHET-LB agar plates inoculated in triplicate with *E. coli* MS75.

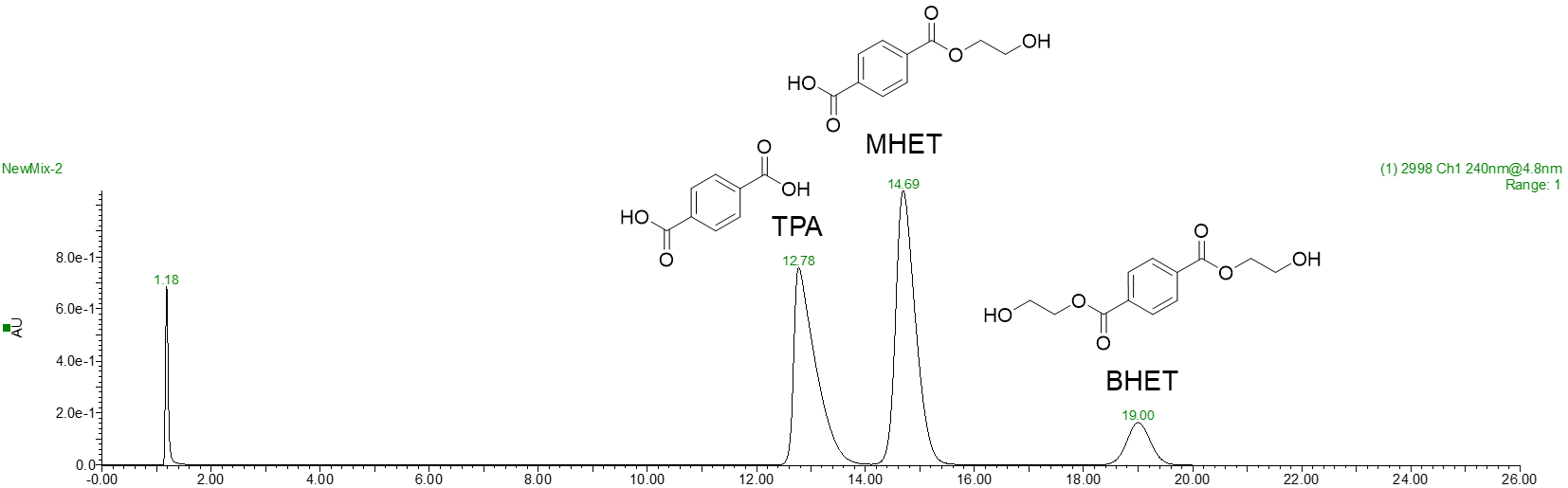

**Figure S13.** Chromatogram obtained by HPLC-PDA showing the peaks related with the standards: BHET (T_R_= 19 min), MHET (T_R_= 14.69 min), and TPA (T_R_= 12.78 min).

AU

| Concentration (mM) | BHET |
| --- | --- |
| 0 | 0 |
| 0.1 | 143596 |
| 0.25 | 348990 |
| 0.5 | 693488 |
| 0.75 | 1038830 |
| 1 | 1337394 |

AU

| Concentration (mM) | MHET |
| --- | --- |
| 1 | 1793608 |
| 0.75 | 1394019 |
| 0.5 | 974349 |
| 0.25 | 499800 |
| 0.1 | 198361 |
| 0 | 0 |

AU

| Concentration (mM) | TPA |
| --- | --- |
| 0.25 | 399359 |
| 0.1 | 180232 |
| 0.05 | 91993 |
| 0 | 530 |

**Figure S14.** Calibration curve of BHET, MHET and TPA at 240 nm.

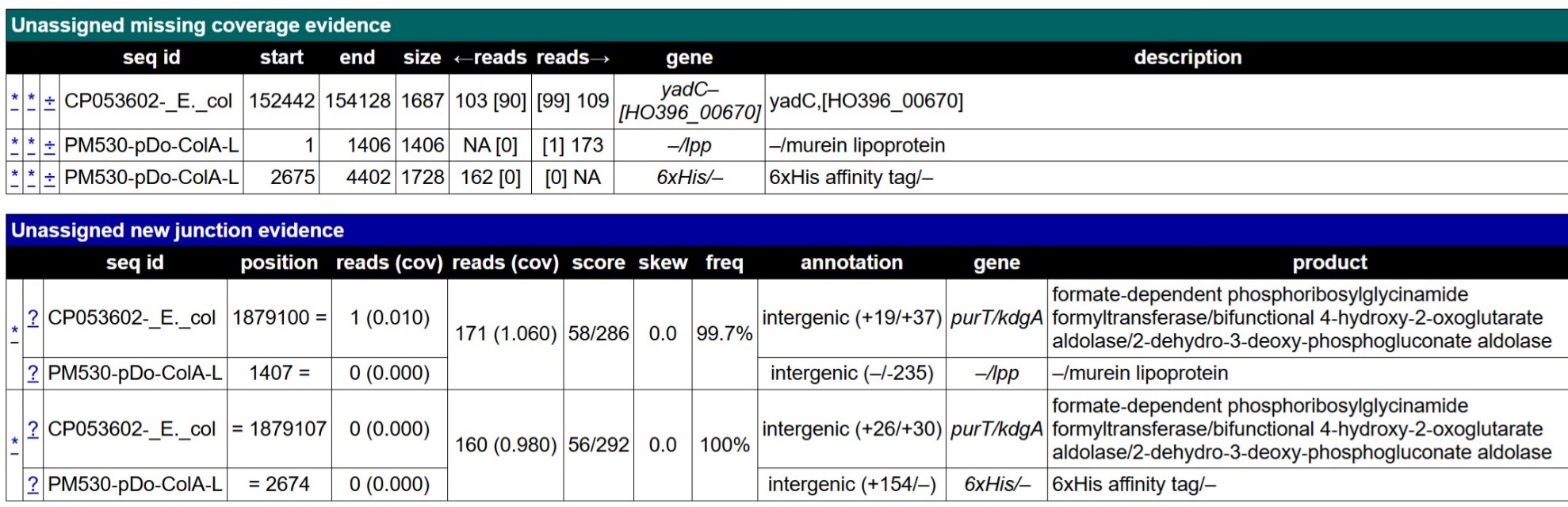

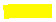

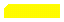

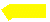

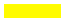

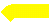

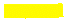

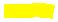

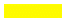

**Figure S15.** Unassigned missing coverage summary and new junction evidence summary after running Breseq. The short-read next generation sequencing results were compared with *E. coli* BL21 (DE3) genome sequence (Ac# CP053602) as well as PM530 donor vector sequence. The highlighted rows show the position of LPP-Cys-PETase cassette insertion into the genome downstream of *kdgA* gene. The position of the junctions at cargo DNA in the donor plasmid (PM530) has also been highlighted.
